# A particle size threshold governs diffusion and segregation of PAR-3 during cell polarization

**DOI:** 10.1101/2021.06.07.447386

**Authors:** Yiran Chang, Daniel J. Dickinson

## Abstract

Regulation of subcellular components’ localization and motion is a critical theme in cell biology. Cells use the actomyosin cortex to regulate protein distribution on the plasma membrane, but the interplay between membrane binding, cortical movements and protein distribution remains poorly understood. In a polarizing one-cell stage *Caenorhabditis elegans* embryo, actomyosin flows transport PAR protein complexes into an anterior cortical domain to establish the anterior-posterior axis of the animal. Oligomerization of a key scaffold protein, PAR-3, is required for aPAR cortical localization and segregation. Although PAR-3 oligomerization is essential for polarization, it remains unclear how oligomer size contributes to aPAR segregation because PAR-3 oligomers are a heterogeneous population of many different sizes. To address this question, we engineered PAR-3 to defined sizes. We report that PAR-3 trimers are necessary and sufficient for PAR-3 function during polarization and later embryo development, while larger PAR-3 clusters are dispensable. Quantitative analysis of PAR-3 diffusion showed that PAR-3 clusters with three or more subunits are transported by frictional drag and experience extensive collisions with the actomyosin cortex. Our study provides a quantitative model for size-dependent protein transportation of membrane proteins by cortical flow.

## INTRODUCTION

Subcellular components must be properly localized for normal cellular function. Cells have evolved a variety of mechanisms to position subcellular components as large as organelles and as small as single mRNA and protein molecules. The actomyosin cytoskeleton is actively involved in many of these mechanisms. For instance, myosin motor proteins transport large cargos, such as vesicles, by ‘walking’ on actin filaments (Mehta, 2001), and chromosome congression in starfish oocytes is accomplished by chromosome trapping within the contracting filamentous actin meshwork (Lénárt et al., 2005). Bulk cytoplasmic and cortical flows, triggered by non-muscle myosin dependent cytoskeleton contraction, are also responsible for the advective transport of a variety of subcellular components in a wide range of organisms, ranging from the circulation of chloroplasts in Elodea leaf cells (Allen and Allen, 1978) to the segregation of PAR proteins in *Caenorhabditis elegans* embryos (Lang and Munro, 2017). However, it has remained unclear how cortical flows can overcome the random motion caused by Brownian diffusion in order to transport macromolecular complexes over large distances.

The *C.elegans* zygote has been widely used to study protein segregation by actomyosin flows due to its relatively large size, optical transparency and the genetic tools that are available. During the first cell cycle of *C. elegans* embryonic development, distinct groups of PAR polarity proteins are segregated to anterior and posterior poles of the cell cortex, leading to asymmetric cell division which is essential for subsequent development. The anterior PAR complex (aPAR) consists of the central oligomeric scaffold protein PAR-3, along with atypical protein kinase C (aPKC/PKC-3) and its cofactor PAR-6. These proteins are distributed uniformly throughout the entire embryo cortex before polarization starts (Cuenca et al., 2003; Lang and Munro, 2017). Upon symmetry breaking triggered by sperm entry, a gradient in actomyosin contractility along the anterior-posterior axis is introduced, and anterior directed cortical and cytoplasmic flows are triggered as a result (Mayer et al., 2010; Munro et al., 2004). aPAR complexes are carried toward the anterior pole by this cortical flow and become enriched at the anterior cell cortex. Once localized, aPKC phosphorylates substrates essential for polarized cell behavior (Hong, 2018; Lang and Munro, 2017).

Multiple studies using different methods have shown that cortical flow is responsible for aPAR segregation in *C. elegans* zygotes. Munro et al. and Shelton et al. demonstrated that cortical flow is required for aPARs segregation by eliminating cortical flow with myosin light chain (MLC-4) RNAi, which results in PAR polarity defect (Munro et al., 2004; Shelton et al., 1999).

Using modeling approaches, Grill and colleagues have shown that advective flow can be sufficient to explain PAR protein partitioning *in silico (Goehring et al., 2011a; Gross et al., 2019)*. Finally, direct induction of cytoplasmic flow by localized laser-induced heating was found to be sufficient to displace the aPAR cortical domain *in vivo* (Mittasch et al., 2018).

However, these studies have not addressed how cortical flows physically transport PAR proteins. The cell cortex consists of actin and myosin filaments as well as surrounding protein and water molecules, but models generally treat the entire cortex as a film of active fluid (Goehring et al., 2011a; Gross et al., 2019; Mayer et al., 2010). Therefore, it remains unclear whether cortical PAR proteins are pushed along the membrane by physical association with actin filaments or, instead, transported through viscous forces generated by cytoplasmic flow.

Clustering of aPAR, which occurs due to PAR-3 oligomerization, is critical for aPAR localization across a range of organisms and cell types (Benton and Johnston, 2003; Dickinson et al., 2017; Mizuno et al., 2003; Rodriguez et al., 2017; Sailer et al., 2015; Wang et al., 2017a). Using CRISPR-induced targeted mutagenesis and live imaging of *C.elegans* embryos, we previously showed that PAR-3 clusters of different oligomer sizes have distinct responses to cortical flow, while mutants in which PAR-3 is strictly monomeric have severe polarity defects and are unable to segregate aPAR (Dickinson et al., 2017). These results hinted at a model in which a large size enables PAR-3 clusters to be transported by cortical flow for proper polarity establishment. However, since PAR-3 oligomers have a wide range of sizes *in vivo* (Dickinson et al., 2017; Lang and Munro, 2017; Wang et al., 2017b), it remains unclear how oligomer size contributes to aPAR segregation.

Here, we address this question by performing a quantitative analysis of oligomerization-dependent protein segregation by actomyosin cortical flow. By dissecting out the heterogeneous PAR-3 population through engineered PAR-3 variants that form oligomers of defined sizes, we reveal that PAR-3 clusters physically collide with actomyosin cortex, and segregate to the anterior cortical domain as a result of viscous friction and collisions with actin rather than a direct, stable association with actomyosin. These results provide fundamental insights into the mechanisms of polarization and of protein transport by cortical flows.

## RESULTS

### Engineered PAR-3 trimers are sufficient for aPAR segregation and normal development

Wild-type PAR-3 exists *in vivo* as a heterogeneous population of oligomers of different sizes, ranging from monomers to >15-mers (Dickinson et al., 2017). In previous work, we disrupted the PAR-3 oligomerization domain by introducing charge-reversal mutations at three key positions in the endogenous *par-3* gene using CRISPR. The resulting PAR-3(RRKEEE) monomeric mutant protein (here referred to as PAR-3*) does not localize stably to the cell membrane or segregate to the anterior domain (Dickinson et al., 2017) (see Figures 1C and Figure S1E). In the same study, we also showed that the brightest 25% of PAR-3 clusters segregated efficiently on the cell cortex in living embryos, while the dimmest 25% moved shorter distances in an apparently diffusive manner. Therefore, large PAR-3 clusters, formed by oligomerization, were presumed to play an important functional role in polarity establishment. However, the exact relationship between PAR-3 cluster size and segregation remained unclear. To dissect the behavior of the heterogenous PAR-3 population and to determine how differently-sized PAR-3 clusters are transported by cortical flow, we engineered PAR-3 oligomers to controlled sizes.

**Figure 1.**
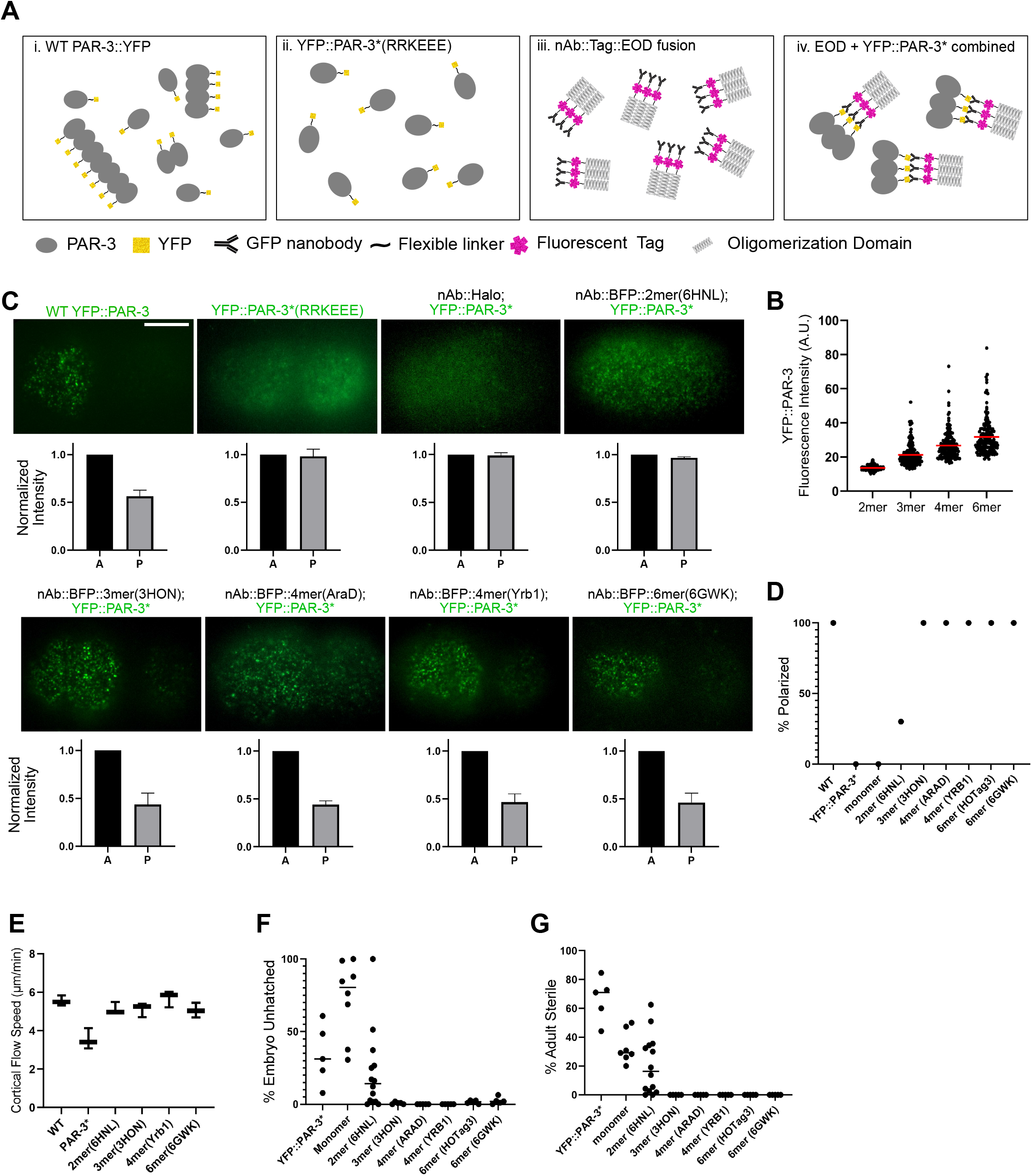
Engineered PAR-3 trimers are sufficient for aPAR segregation and normal development. (A) Illustration of our strategy to engineer PAR-3 to specific sizes. i. During polarity establishment, PAR-3 clusters exist as a heterogeneous population of various sizes. ii. PAR-3 monomers in a mutant embryo in which the PAR-3 oligomerization domain is disrupted. iii. Transgenic EOD constructs contain the EOD (3mer is shown as an example), a fluorescent label and a GFP-binidng nanobody. iv. PAR-3 of controlled oligomer size was generated by combining YFP::PAR-3*(RRKEEE) and EOD constructs. (B) YFP fluorescence intensity of the foci on the cortex of YFP::PAR-3*;nAB::BFP::EOD embryos dissected from *mlc-4* RNAi treated worms. *mlc-4* RNAi was used for these measurements to prevent crowding of PAR-3 clusters into the anterior domain, which makes accurately measuring intensity more difficult. Data shows only clusters that are stabilized on the cortex for more than 10 frames (0.5 seconds), because these are the particle tracks we used in the quantitative analysis below. We obtained equivalent results with or without the track length filter (Figure S1F). Red bars indicate means. (C) Upper panels: Live images of cortical YFP::PAR-3* in the indicated EOD strains. Scale bar: 10 μm. Lower panels: Quantification of fluorescence intensity in posterior area to anterior area. n = 3. (D) Percentage of embryos from each EOD strains that display effective PAR-3 polarization. n = 10. (E) Cortical flow speed measured from DIC movies of live embryos dissected from wild-type, YFP::PAR-3*(RRKEEE) and YFP::PAR-3*;nAB::BFP::EOD strains. n = 3. Horizontal bars indicate means. (F) Embryo lethality of each EOD strain. “Monomer” is a control construct comprising the nanobody and fluorescent tag but no EOD. Horizontal bars indicate means. (G) Adult sterility of each EOD strain. “Monomer” is a control construct comprising the nanobody and fluorescent tag but no EOD. Horizontal bars indicate means.

We tested a handful of protein domains that have been reported to autonomously organize into oligomers of defined sizes *in vitro* (Figure S1A-B) (Bolten et al., 2016; Boudko et al., 2009; Büttner et al., 2012; Chik et al., 2019; Drulyte et al., 2019; Huang et al., 2014; Lee et al., 1968; Li et al., 2019; Luo et al., 2001; Parsons et al., 2002; Santiago-Frangos et al., 2019; Sun et al., 2014; Thomson et al., 2014; Veesler et al., 2010). Each oligomerization domain was tagged with a fluorescent protein and expressed in *C.elegans,* and the size of each oligomer protein complex from the transgenic zygotes is examined using single-cell, single-molecule pull-down (sc-SiMPull) followed by photobleaching step counting (Dickinson et al., 2017) (Figure S1B). We identified dimer, trimer, tetramer and hexamer protein domains that formed oligomers of the expected sizes *in vivo*.

To generate PAR-3 oligomers of defined sizes, we adopted a strategy in which PAR-3 is linked to one of these “extra oligomerization domains” (EODs) via a nanobody. We generated transgenic constructs consisting of a nanobody that binds to GFP/YFP (Wang et al., 2017a), a fluorescent BFP or HaloTag, and an EOD (Figure 1A iii). We verified that the nanobody successfully bound to GFP/YFP in *C. elegans* lysates *in vitro* (Figure S1D) and in live embryos *in vivo* (see below). To generate engineered strains with PAR-3 oligomers of controlled sizes, we crossed each nAb::EOD construct to a strain carrying an endogenous YFP-tagged PAR-3 monomer allele (YFP::PAR-3*, which has the RRKEEE charge reversal mutations identified previously (Dickinson et al., 2017)) (Figure 1A). We measured the YFP intensity of these induced foci on the cortex and confirmed that they exhibited the expected trend of increasing sizes (Figure 1B).

Strikingly, our 3mer, 4mer and 6mer nAb::EOD constructs fully rescued the polarity defects in the parental YFP::PAR-3* monomeric strain. Figure S1E illustrates this rescue using one of the 6mer constructs (HOTag3) as an example. Neither the 6mer(HOTag3) EOD construct nor YFP::PAR-3* monomer by itself localized to the cortex or adopted a polarized distribution, but when the 6mer(HOTag3) EOD construct and YFP::PAR-3* were present together, the two proteins colocalized in clusters that segregated to the anterior half of the zygote (Figure S1E). The engineered 3mer(3HON), another 6mer(6GW3) and two different 4mers(AraD and Yrb1) also formed clusters that segregated to the anterior (Figures 1C-D and Movie S1).

A negative control strain without any EOD, nAb::Halo;YFP::PAR-3*, closely resembled the YFP::PAR-3* mutant: we observed only very weak membrane binding, with only a few particles visiting the membrane for a very short amount of time and without forming a polarized anterior domain (Figure 1C and Movie S1). The engineered 2mer (6HNL) PAR-3 exhibited an intermediate phenotype: PAR-3 dimers localized to the membrane with shorter membrane binding lifetime (Movie S1; see Figure 5B-C, below, for quantification). A minority of 2mer embryos (4/14) showed some ability to polarize, but most embryos (10/14) were not polarized effectively (Figure 1D and S1G). These data indicate that a threshold size of three PAR-3 molecules per cluster is necessary and sufficient for robust polarity establishment in the *C. elegans* zygote.

Importantly, the difference in polarization between 2mer and larger EODs is not due to differences in actomyosin cortical flow speed (Figure 1E). Previous studies showed that cortical flow speed was significantly reduced in PAR-3 monomeric mutant strains, although this reduction in cortical flow was not severe enough to account for loss of polarity in PAR-3 monomer strains (Rodriguez et al., 2017). To ensure that the differences in EOD strain polarization were not due to differences in cortical flow, we aquired DIC movies of live EOD emrbyos and measured the cortical flow speed.We confirmed the slower cortical flow speed seen earlier in PAR-3 monomeric mutants (Rodriguez et al., 2017), but all of the EOD strains, including 2mer, showed cortical flow speeds similar to wild-type (Figure 1E). We conclude that the difference in polarization between 2mer and larger EODs is due to factors other than cortical flow speed.

On the phenotypic level, strains carrying the monomeric PAR-3* mutation exhibit partially penetrant embryonic lethality and adult sterility (Dickinson et al., 2017). Remarkably, these phenotypes were almost fully rescued by the 3mer, 4mer or 6mer EOD::nAb constructs, and slightly rescued by the 2mer construct (Figure 1F-G). We did observe some mild phenotypes in two of the individual engineered strains: the PAR-3*; 6mer(HOTag3) strain had slow larval growth and maturation, and a few embryos (<10%) of the PAR-3*;4mer(AraD) strain exhibited cytokinesis defects. These phenotypes are not present in other stains and have no obvious relationship to cell polarity in the zygote, and we presume that they are artifacts of expressing these specific EODs. We excluded these two strains from the analysis that follows. Additionally, while we did not observe plate-level phenotypes in the 4mer(Yrb1) strain, we noticed that embryos dissected from this strain were more delicate; they more often died or took a longer time to polarize during fluorescence imaging. Despite these caveats of specific engineered EOD strains, the recovery of PAR-3 localization, zygote polarity, embryonic lethality and adult sterility indicate that oligomers as small as trimers are sufficient for PAR-3 function in *C. elegans*.

### PAR-3 trimers and larger oligomers undergo directed motion due to cortical flow, while PAR-3 dimers do not

We were intrigued by the observation that engineered PAR-3 3mers were sufficient to enable cell polarization, but 2mers were not. To attempt to explain why cortical flow is effective on 3mers but not on 2mers, and to better define the movement of PAR-3 on the cortex, we analyzed the diffusive behavior of individual engineered PAR-3 clusters. We imaged one-cell embryos of each EOD strain during polarity establishment using TIRF microscopy at a high frame rate (20 frames/second) to visualize the diffusion of clusters on the cortex, and then computationally segmented and tracked individual particles (Figure 2A). To characterize and compare the motion of different-sized clusters, we performed a Mean Squared Displacement (MSD) analysis. The diffusion of particles on a 2D surface, such as the cell membrane, can be described as:

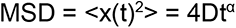

**Figure 2.**
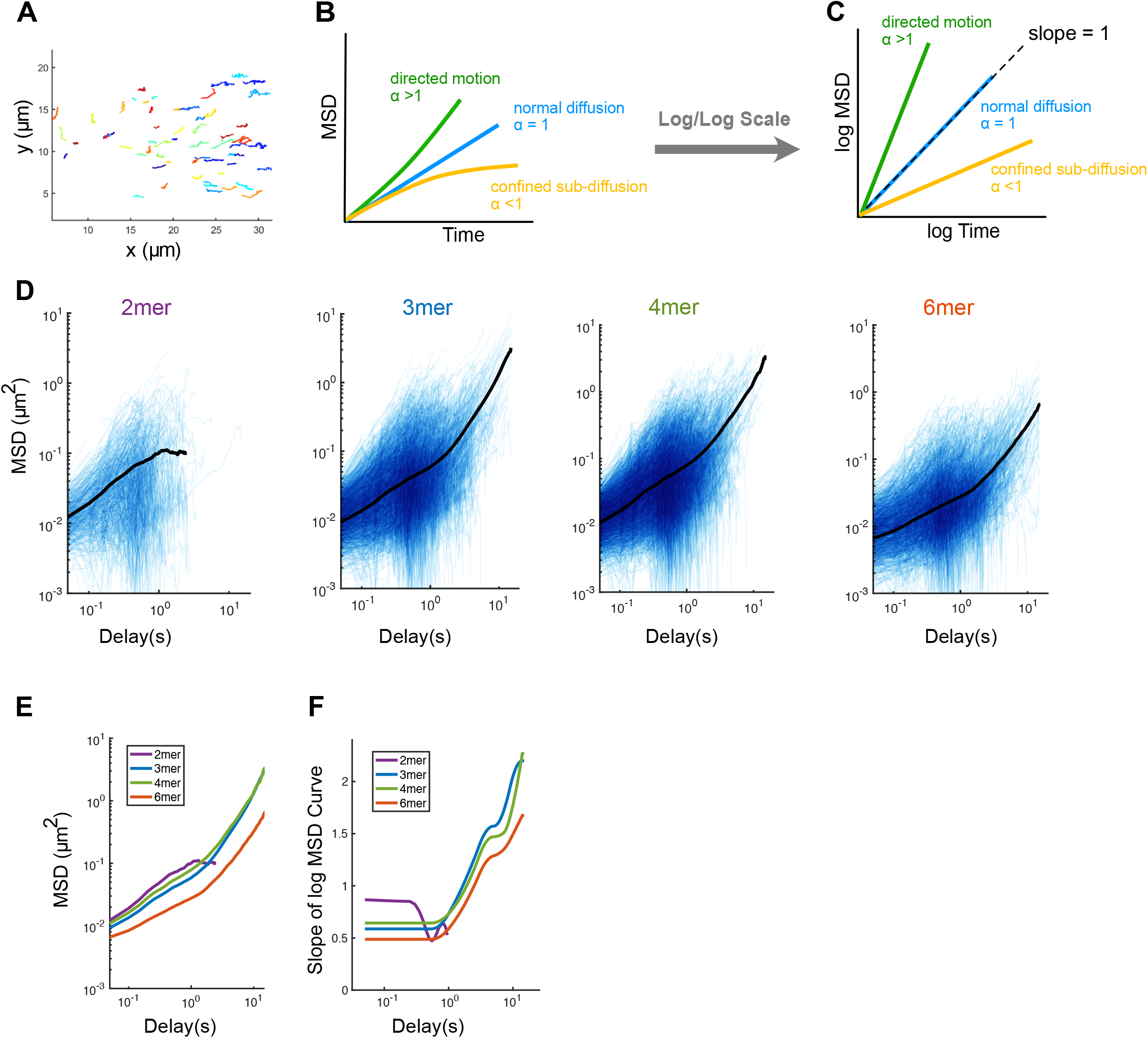
PAR-3 trimers and larger oligomers undergo directed motion due to cortical flow, while PAR-3 dimers do not. (A) Tracks of engineered PAR-3 trimers imaged at the cortex during polarity establishment. Anterior to the left. Different colors are only used for better visualization. t = 15 seconds. (B) Illustration of the interpretation of anomalous parameters. (C) Illustration of the interpretation of the MSD curve on log/log scales. (D) log/log scale MSD curves for 2mer, 3mer, 4mer and 6mer, each curve describing the motion of a single PAR-3 cluster. For each plot, data were acquired and pooled from 3 embryos. (E) Comparison of the averaged MSD curves for 2mer, 3mer, 4mer and 6mer. (F) Slopes of the averaged MSD curves for 2mer, 3mer, 4mer and 6mer, estimated by fitting a smoothing spline to each curve and then taking its derivative.

Where D is the diffusion coefficient and α is the anomalous diffusion parameter. α = 1 describes normal Brownian diffusion. An α <1 indicates sub-diffusion where the movement of a particle is confined, while α > 1 represents super-diffusion, which indicates directed movement of particles (Figure 2B-C). Therefore, the anomalous parameter provides direct information about the type of motion a particle is experiencing.

The anomalous parameter α can be easily visualized by plotting MSD vs. time on a log/log scale: log-transformed data for diffusing particles are lines with a slope equal to α. For particles undergoing normal Brownian diffusion, these plots are straight lines with a slope of 1. To measure the anomalous diffusion parameters for PAR-3 clusters of different sizes, we plotted MSD on a log/log scale for 2mer(6HNL), 3mer(3HON), 4mer(Yrb1), and 6mer(6GW3) EOD strains (Figure 2D). We generated average MSD curves (Figure 2E) and calculated the slope of each log/log MSD curve by taking the derivative of a smoothing spline fit to the data (Figure 2F). The 4mer(AraD) and 6mer(HOTag3) EOD strains were not included in this analysis due to the mild, non-polarity related phenotypes noted above.

This MSD analysis revealed non-Brownian diffusive behaviors of PAR-3 oligomers, which are distinct between engineered dimer and other larger engineered oligomers. The log-log MSD curves for 3mers, 4mers and 6mers are not straight lines (Figure 2D-E), which indicates that these clusters undergo different types of motion on different timescales. On the shorter time scale (<1s), the anomalous parameter falls below 1 for clusters of all sizes, which suggests a confined movement, presumably due to the physical interaction with the cortical actomyosin network. However, 2mers had a slope closer to 1 compared to larger clusters (Figure 2F), indicating that their movement is more diffusive than that of larger clusters. On longer time scales (>1s), larger oligomers display anomalous parameters that increase to greater than 1, which suggests a directed motion under the effect of cortical flow (Figure 2D-F). In contrast, 2mers did not exhibit directed motion but instead dissociated from the cortex on longer timescales (Figure 2D).

The observed difference between the dimers and larger oligomers is consistent with the polarity and embryo lethality phenotypes we observed in the 2mer strain (Figure 1). We conclude that during polarization, movement of PAR-3 2mers on the cortex is dominated by weakly confined random diffusion, while 3mers, 4mers and 6mers undergo directed movement under cortical flow at longer time scales.

### PAR-3 clusters and cortical actin move in tandem but are not physically associated

To address the physical explanation for the non-Brownian diffusive behavior of larger PAR-3 clusters, we examined the interaction between PAR-3 and the actomyosin cortex. Although the actomyosin cortex and PAR-3 clusters move in tandem towards the anterior pole, there is no known binding interaction between PAR-3 and F-actin. The relationship between PAR-3 and actomyosin cortex has been directly visualized by observing labeled myosin II (NMY-2) (Movie S2) (Dickinson et al., 2017), but F-actin and PAR-3 have not been visualized together in living embryos to our knowledge. Therefore, we constructed a strain carrying endogenously tagged mScarlet::PAR-3 and a transgenic GFP::utrophin reporter that binds to F-actin (Tse et al., 2012) by endogenously tagging PAR-3 with mScarlet-I in a GFP::utrophin background. We first imaged zygotes from this strain at high speed (3seconds/frame) using TIRF microscopy (Figure S2A and Movie S3). In agreement with previous studies, live fluorescence imaging showed that the movement of the actin cortex and of PAR-3 are tightly coupled (Goehring et al., 2011a; Munro et al., 2004) (Figure 3A-B and Movie S3).

**Figure 3.**
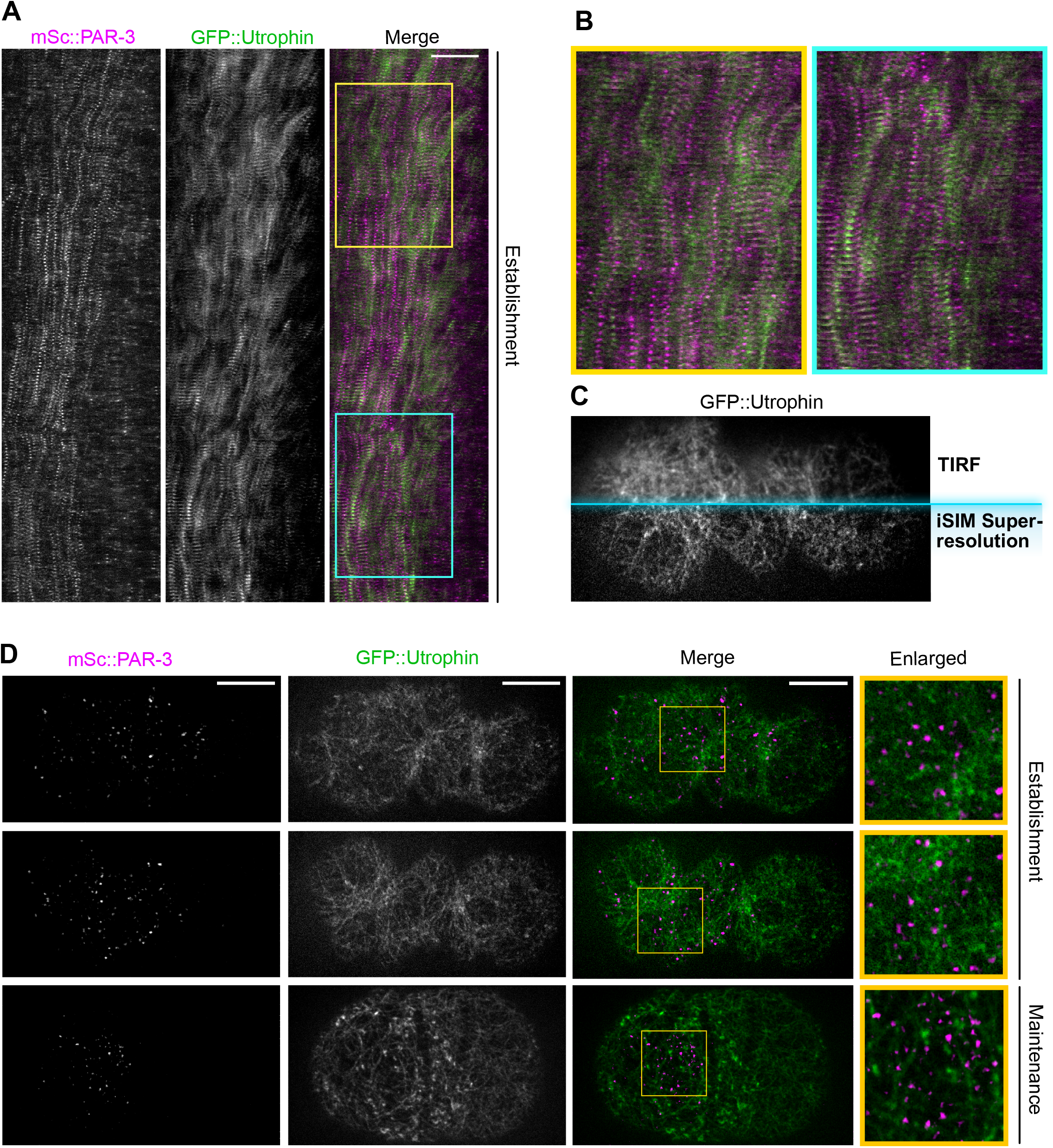
Super-resolution imaging reveals that PAR-3 moves in tandem with the actomyosin cortex despite not being physically associated with actin. (A) Kymograph of cortical mSc::PAR-3 and Utrophin::GFP during the first cell cycle. Imaged using TIRF at 3 seconds/frame. Anterior to the left. Scale bar = 10µm. Yellow/Blue boxes: region enlarged in (B). (B) Enlarged sections of kymograph (boxes) in (A). (C) The comparison between TIRF imaging and high resolution iSIM imaging. A mSc::PAR-3;GFP::UTRO zygote at polarization stage is imaged and only GFP::Utrophin channel is shown. (D) Maximum projected Z stacks of images of mSc::PAR-3;GFP::Utrophin zygote, imaged using super-resolution iSIM imaging centered at the cortical region . Anterior to the left. Scale bar = 10µm. Yellow box: region enlarged in right column.

To quantify the correlation between PAR-3 segregation and actin flow, we selected 2 time points during polarity establishment: one when the cortex was actively contracting towards the anterior pole and one when the cortex was temporarily oscillating and moving towards the future dorsal side. We quantified the movements using particle image velocimetry (PIV) and compared the vector fields for PAR-3 and F-actin using Pearson’s correlation (Figure S2B–C). As expected, we observed a positive correlation between PAR-3 movement and F-actin movement (Pearson’s correlation R = 0.70 for Δt1, R = 0.59 for Δt2), indicating that PAR-3 and the actomyosin cortex move in tandem. Nevertheless, PAR-3 and actin did not colocalize or overlap with each other in TIRF images (Pearson’s correlation R = 0.067 ± 0.010; Figure 3A-B).

Although we did not observe an obvious association between PAR-3 and actin in TIRF images, it remained possible that contacts between actin and PAR-3 might have been overlooked due to the density of the F-actin network and the diffraction-limited resolution of our TIRF images.

Therefore, to better visualize the interaction between PAR-3 and the actin network, we performed super-resolution imaging of the cortex using Instant Structured Illumination (iSIM) (Figure 3C). We found that the PAR-3 clusters appeared as point sources even in super-resolution images, suggesting that even the largest clusters are smaller than the ∼170 nm resolution of iSIM imaging (Figure 3D). In contrast, individual actin branches and pores in the actomyosin cortex were clearly visible (Figure 3D). These observations reveal important information about the scale of sizes: PAR-3 clusters are much smaller than the gaps in the actomyosin cortex. Furthermore, PAR-3 clusters were frequently found within the pores of the cortex, not associated with individual filaments (Figure 3D).

In a previous study, Sailer et al. found that during polarity maintenance, after polarizing cortical flows have ceased, wild-type PAR-3 clusters undergo weakly confined diffusion at the cortex, especially on timescales longer than 1s (Sailer et al., 2015). Treatment with an actin depolymerizing drug eliminated this confined behavior, suggesting confinement was due to interactions between PAR-3 and F-actin (Sailer et al., 2015). These results are consistent with a “corralling” effect, where diffusion of PAR-3 clusters is restrained by collisions with a dynamic actin network. Our finding that PAR-3 clusters are located within pores in the actin network (Figure 3D) supports this idea: although we find no evidence for a direct association between PAR-3 and actin, PAR-3 clusters that diffuse within pores of the actin cortex would be expected to exhibit confined motion due to collisions with actin, especially on longer timescales.

To further explore this idea, we sought to directly test whether the larger PAR-3 oligomers are confined within the actomyosin meshwork of the cortex during polarity establishment. We repeated the imaging and MSD analysis in our EOD strains after eliminating cortical flow by depleting myosin light chain (*mlc-4* RNAi), which allowed us to observe the behavior of PAR-3 clusters in the absence of cortical flow. Importantly, *mlc-4* RNAi eliminated cortical flow but did not grossly disrupt the organization of the actin cortex (Figure 4A). As expected, *mlc-4* depletion eliminated directed movement of larger PAR-3 clusters during polarity establishment (Figure 4B-D; compare to Figure 2D-F). Instead, PAR-3 clusters exhibited sub-diffusive movement that became progressively more confined on longer timescales (Figure 4C-D), which is nearly identical to the behavior previously reported for wild-type PAR-3 clusters during polarity maintenance (Sailer et al., 2015). We noted that despite exhibiting sub-diffusive behavior,

**Figure 4.**
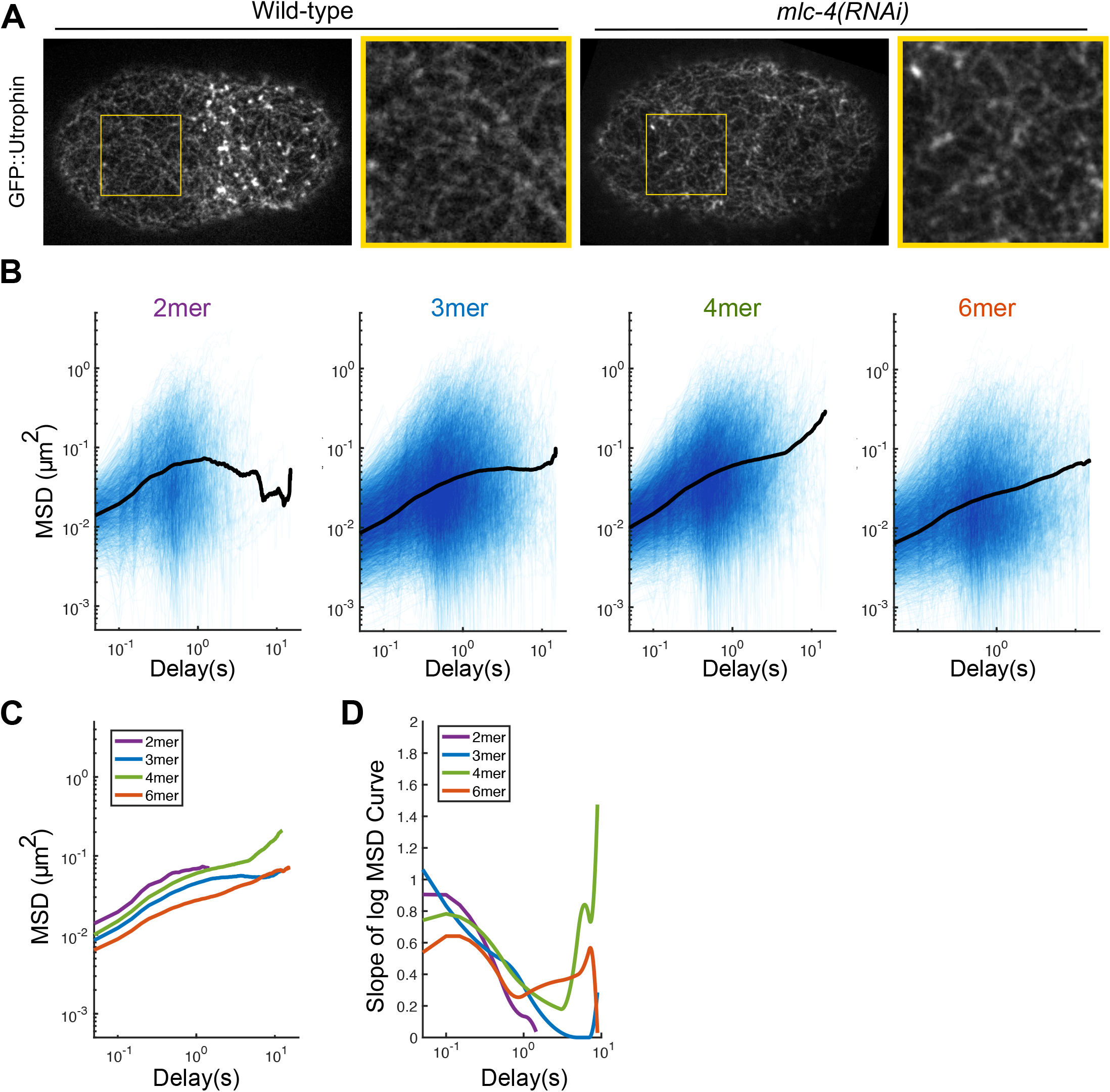
MSD Analysis with cortical flow eliminated reveals PAR-3 clusters are loosely confined by the transient physical interactions with the actomyosin cortex. (A) Super-resolution iSIM images of GFP::Utrophin zygotes at the pronuclei meeting stage, with or without *mlc-4* RNAi. The structure of the actin network is intact under *mlc-4* RNAi. Yellow box: region enlarged to the right of each image. (B) log/log scale MSD curves for 2mer, 3mer, 4mer and 6mer with *mlc-4* RNAi treatment, each curve describing the motion of a single PAR-3 cluster. For each plot, data was acquired and pooled from 3 embryos. (C) Comparison of the averaged MSD curves for 2mer, 3mer, 4mer and 6mer. Curves were trimmed to remove the noisy tails at long timescales that result from only a few particles. (D) Slopes of the averaged MSD curves for 2mer, 3mer, 4mer and 6mer, estimated by fitting a smoothing spline to each curve and then taking its derivative.

PAR-3 clusters remained mobile even under *mlc-4* RNAi conditions. Together with our imaging results (Figure 3), these data rule out models in which PAR-3 is directly bound to F-actin or tightly encased in a dense actin meshwork. Instead, our results suggest that PAR-3 clusters diffuse within pockets in the actin cortex and are partially confined via collisions with F-actin.

An additional observation from these experiments was that engineered PAR-3 clusters, regardless of size, moved in a sub-diffusive fashion when *mlc-4* was depleted. There was no clear difference between 2mers and larger oligomers in these experiments. Therefore, the failure of the 2mer construct to rescue polarity (Figure 1) cannot be explained by differences in confinement or corralling of PAR-3 2mers compared to larger oligomers. We therefore explored alternatively possible explanations for the inability of 2mers to support normal polarity establishment.

### Differences in diffusivity and membrane lifetime together can account for the different behaviors of 2mer and larger oligomers

Up to this point, we have identified differences in diffusive behavior between 2mers and larger oligomers, but how these observations are linked to the different polarization effectiveness in EOD strains remains unknown. To attempt to quantitatively explain the difference in polarization between 2mers and larger oligomers, we first measured the diffusion coefficients and membrane binding lifetimes of engineered PAR-3 oligomers. To estimate the diffusion coefficients, we fit the first six time steps of each averaged MSD curve, corresponding to timescales before the motion became significantly confined, to the diffusion equation (MSD = 4Dt) (Figure S3A). Our measured diffusion coefficients (Figure 5A) were much smaller than those observed for another anterior PAR protein, PAR-6, in bulk FRAP experiments (Goehring et al., 2011b) but were similar to those of a slowly diffusing sub-population of PAR-6 particles that was likely associated with PAR-3 (Robin et al., 2014).

**Figure 5.**
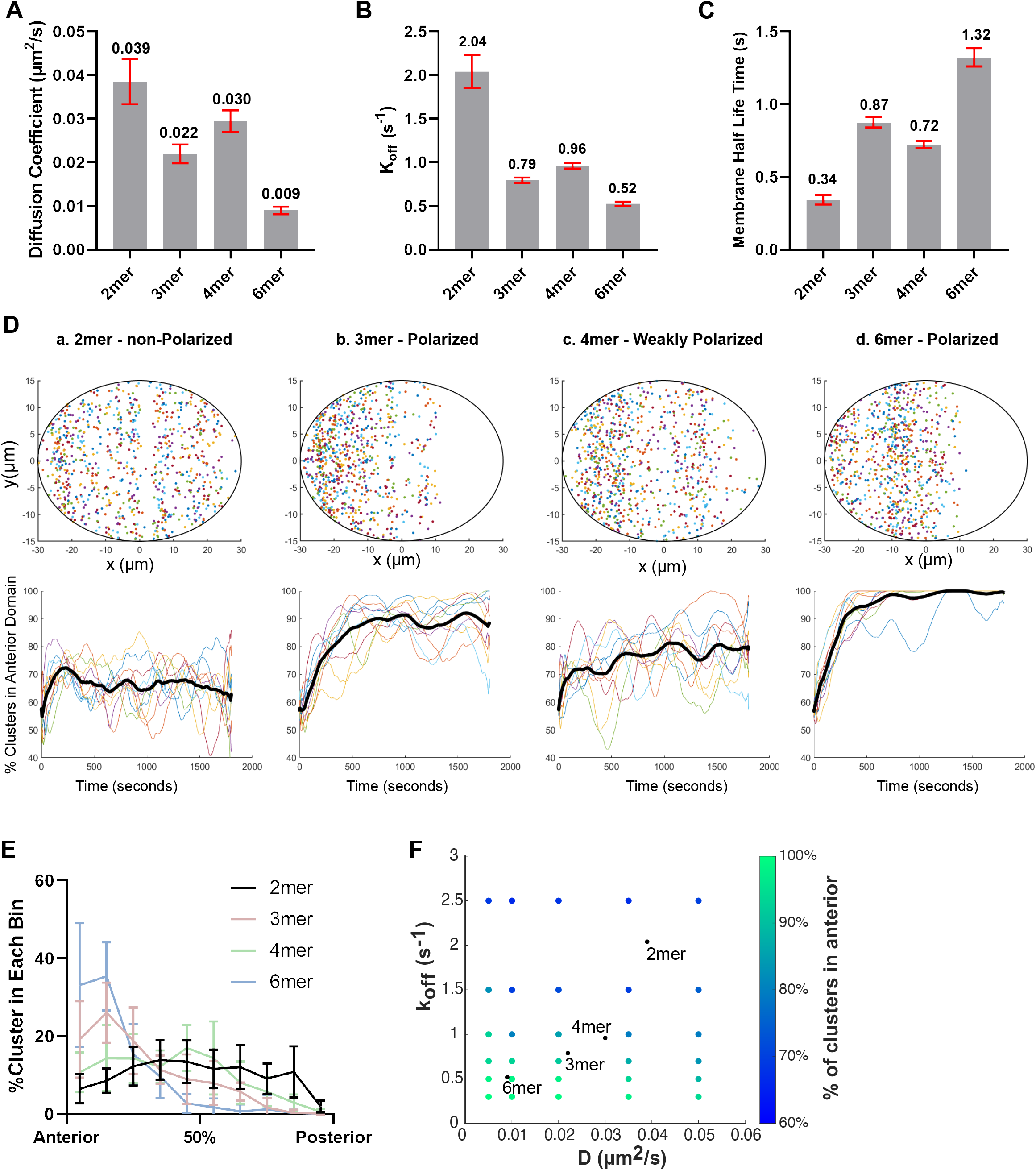
Simulations reveal that the different behaviors of dimers and larger oligomers can be explained by combining diffusion and membrane binding. (A) The diffusion coefficient in engineered PAR-3 strains (wild-type background). Red bars indicated 95%CI. Data points pooled from 3 embryos for each strain. n = 4237, 10587, 12619, 6724 particles for 2mer, 3mer, 4mer, 6mer, respectively. (B) The k_off_ in engineered PAR-3 strains (wild-type background). Red bars indicated 95%CI. Data points pooled from 3 embryos for each strain. n = 4237, 10587, 12619, 6724 particles for 2mer, 3mer, 4mer, 6mer, respectively. (C) The membrane binding half-time of engineered PAR-3 strains. with 95% CI. Red bars indicated 95%CI. Data points pooled from 3 embryos for each strain. n = 4237, 10587, 12619, 6724 particles for 2mer, 3mer, 4mer, 6mer, respectively. (D) Upper panels: final cluster positions after simulating particle movement for 15 minutes. Elipse: boundary of simulated embryo. x and y units: μm. Lower panels: percentage of particles in the anterior domain over time in simulated embryos. Each line describes an independent repeat of simulation. Black line: average. (E) The number of clusters in each 6μm wide bin along the AP axis in simulated EOD embryos. n = 10 for each set of conditions. (F) Dependence of the simulation outcomes on parameter values. Each point represents the average results of 10 simulation runs. The parameter values used in (D) and (E) are indicated by black points.

To quantify membrane lifetime, we calculated the particle disappearance probability per imaging frame in each EOD strains, based on the distributions of particle track lengths (see Methods).

Then, using the known imaging frame rate, we converted the particle disappearance probability to a membrane unbinding rate (Figure 5B) and membrane half life time (Figure 5C). A caveat of these estimates is that clusters can disappear due to photobleaching in addition to membrane unbinding, resulting in an overestimation of the actual k_off_. To evaluate the severity of photobleaching in our experiments, we examined fluorescence intensity over time in our 6mer dataset. Since 6mers have the longest membrane lifetime (and therefore reside in the TIRF excitation field for the longest), and can undergo significant photobleaching without disappearing entirely, we would expect their intensity to decrease over the length of a single particle track if appreciable photobleaching were occurring. However, we found that the fluorescence intensity of most 6mers remained constant over the length of our observation; the median 6mer was 94% as bright during the last 10 frames before it disappeared as during the first 10 frames of its observation (Figure S3B). Therefore, photobleaching makes only a minor contribution to our estimates of k_off_ and membrane lifetime.

We observed trends of decreasing diffusivity (Figure 5A) and increasing membrane lifetime (Figure 5B-C) as oligomer size increased. We therefore hypothesized that the slower diffusion and reduced random motion of larger PAR-3 oligomers, together with sufficient membrane dwell time to experience cortical flow, would allow larger oligomers to be efficiently segregated towards the anterior.

To test this hypothesis, we computationally simulated the movement on the plasma membrane of PAR-3 clusters with different diffusivities and membrane binding lifetimes. Prior studies have developed complex and sophisticated models of PAR protein segregation and mutual antagonism (Goehring et al., 2011a; Sailer et al., 2015; Gross et al., 2019). However, these models either did not include PAR protein clustering (Goehring et al., 2011a; Gross et al., 2019) or did not simulate the process of polarity establishment (Sailer et al., 2015). Here, we took a simpler approach in order to focus on how cortical flow would be expected to segregate clustered proteins of different sizes. Each particle in our simulations underwent a biased random walk due to the combination of Brownian diffusion and cortical flow, and at each time step, particles dissociated from the membrane with a probability calculated from k_off_. We incorporated two additional assumptions, which are based on experimental observations. First, we assumed that the number of PAR-3 clusters at the membrane remains constant over time; that is, for each PAR-3 cluster that dissociates from the membrane, a new one appears. This assumption is consistent with the observation that the amount of PAR-3 at the membrane remains roughly constant throughout polarization (Movie S1). Second, we assumed that new PAR-3 clusters preferentially bind the membrane in the part of the cell where PAR-3 is present. This assumption is justified by observations that posterior PAR proteins occupy the region of the plasma membrane that has been cleared of anterior PARs (Cuenca et al. 2003) and that posterior PAR proteins prevent binding of new PAR-3 clusters to the membrane (Sailer et al. 2015). We implemented this assumption by choosing the locations of newly-binding PAR-3 particles at random from the measured distribution of existing particles at each time step. We note that this second assumption introduces positive feedback to the system, because more PAR-3 particles are added to the regions of the membrane where PAR-3 particles are already enriched. To ensure that this form of feedback was not sufficient to establish polarity on its own, we ran simulations in the absence of cortical flow and confirmed that polarization did not occur (Figure S3D). Then, we ran simulations using the diffusion coefficient and k_off_ measured in each EOD strain.

We quantified the polarization state of each simulated embryo in several different ways. First, we ran each simulation 10 times, and we quantified the percentage of clusters in the anterior domain over time in each simulation. We observed an efficient polarization within the first 10 minutes using parameters measured for 3mers and 6mers. In contrast, using parameters measured for 2mers, the percentage of clusters in the anterior domain remained constant for the duration of the simulations (Figure 5D, lower panels). We also quantified the final cluster distribution along the AP axis by counting the clusters in each 6 μm wide bin along the anterior-posterior axis (Figure 5E). Consistent with previous results, simulated PAR-3 3mers and 6mers were localized in the anterior domain, but simulated 2mers remained evenly distributed. Using the parameters measured for 4mers, we observed polarization that was significant, but weaker than that of simulated 3mers and 6mers. This is consistent with the fact that our measurements of D and k_off_ for 4mers fell outside the trend observed for 2mers, 3mers and 6mers. Although the reason for faster diffusion and weaker membrane binding of 4mers is not clear, it is consistent with the mild phenotype observed for 4mer(YrbI) EOD embryos (see above). Overall, our simulation results indicate that the efficient segregation of larger PAR-3 oligomers can be explained by cooperativity between reduced random motion and membrane binding.

To more generally explore how diffusivity and membrane binding lifetime determine the efficiency of polarity establishment, we systematically tested the effect of different combinations of D and k_off_ spanning the range measured from PAR-3 oligomers (Figure 5F). We found that the effectiveness of polarization, quantified by percentage of clusters in the anterior, decreases as k_off_ and diffusion coefficient increase. Intuitively, for cortical flows to effectively segregate a clustered protein, the cortical flow must dominate the randomization of particle positions that occurs due to diffusion and membrane unbinding/rebinding. Our simulations reveal that slower diffusion and more stable membrane association allows larger PAR-3 clusters to be efficiently polarized by cortical flow.

### Tracking wild-type PAR-3 movement with a novel dual-labeling technique confirms the size threshold in an endogenous setting

Up to this point, we have characterized the diffusive behavior of engineered EOD::PAR-3, which revealed size-dependent polarization behavior but represents a somewhat artificial situation. To test whether the size-dependent diffusion behavior of PAR-3 clusters is also true in endogenous conditions, we sought to measure the diffusive behaviors of a heterogeneous population of wild-type PAR-3 clusters. It is challenging to perform this analysis using wild-type PAR-3 tagged with fluorescent proteins, for two reasons: first, there is an inherent bias towards bright signals during particle tracking, and second, the high density of PAR-3 clusters causes challenges in particle tracking, limiting the ability to accurately assemble long tracks for all but the brightest particles. In our previous work (Dickinson et al., 2017), these challenges prevented us from measuring diffusion parameters of endogenously tagged mNeonGreen::PAR-3.

To solve these issues, we developed a new method that allowed us to monitor the diffusion of PAR-3 clusters in an unbiased way while also estimating their sizes. We labeled endogenous HaloTag::PAR-3 with a mixture of two different HaloTag ligand dyes (Grimm et al., 2015, 2016, 2017) at a defined ratio. Each HaloTag molecule binds to a single dye molecule, so that when a mixture of dyes is used for labeling, a mixed population of labeled Halo::PAR-3 molecules results (Figure 6A). By adjusting the ratio of the two dyes, we achieved conditions in which the less-abundant dye was present at single-molecule levels on the cortex, yielding discrete labels that are ideal for tracking. At the same time, the more-abundant dye reveals the cluster size information (Figure 6A).

**Figure 6.**
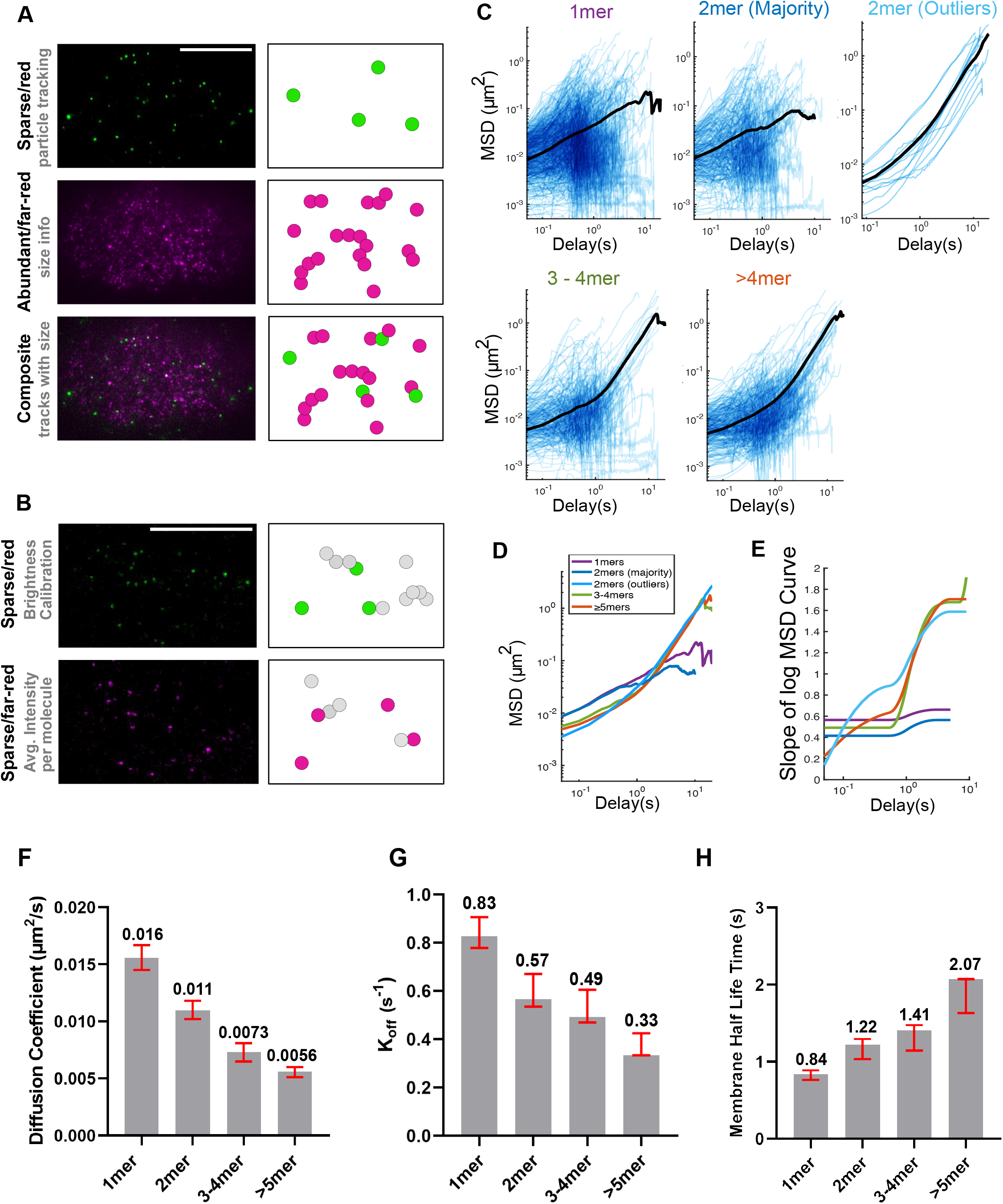
The PAR-3 size threshold in an endogenous setting. (A) TIRF images and cartoon illustrations of the dual-labeling experiment. Magenta represents far-red/abundant channel and green represents red/sparse channel. (B) TIRF images and cartoon illustrations of the calibration double dilution experiment, where the dyes for both channels are diluted to single molecule levels. (C) log/log scale MSD curves for each cluster group binned by estimated size, each curve describing the motion of a single PAR-3 cluster. For each plot, data were acquired and pooled from 5 embryos. (D) Comparison of the averaged MSD curves for each cluster group binned by estimated size. (E) Slopes of the averaged MSD curves for each cluster group binned by estimated size. Slopes were estimated by fitting a smoothing spline to each curve and then taking its derivative. (F) The diffusion coefficient measured in a dual-labeling experiment. Red bars indicated 95%CI. Data points pooled from 3 embryos for each strain. n = 672, 297, 244, 322 particles for 1mers, 2mers, 3-4mers, >5mers, respectively. (G) The k_off_ measured in a dual-labeling experiment. Red bars indicated 95%CI. Data points pooled from 3 embryos for each strain. n = 672, 297, 244, 322 particles for 1mers, 2mers, 3-4mers, >5mers, respectively. (H) The diffusion coefficient measured in a dual-labeling experiment. Red bars indicated 95%CI. Data points pooled from 3 embryos for each strain. n = 672, 297, 244, 322 particles for 1mers, 2mers, 3-4mers, >5mers, respectively.

We imaged dual-labeled Halo::PAR-3 embryos at 20 frames/second on a custom-built TIRF microscope with a dual-view emission path design, allowing simultaneous imaging of both wavelengths. We first analyzed the images of the sparse (red) dye, using the same method as for EOD strains, to perform the MSD analysis. Then, we found the relative size of each particle by fetching the fluorescence intensity of each particle in the abundant (far-red) channel (Figure 6A). To convert this relative size, in the form of fluorescence intensity, into an estimate of molecules per cluster, we adopted a calibration procedure using the fluorescence intensities of single sparse (red) molecules as an internal standard. In brief, we diluted both HaloTag ligand dyes to single-molecule levels and collected images under identical imaging conditions as those used for the diffusion analysis above; this allowed us to measure the brightness of single far-red dye molecules, which we then used to estimate the number of HaloTag::PAR-3 molecules in each cluster in our sparse/abundant experiments (Figure 6B, S4 and Methods).

Tracking single endogenous PAR-3 clusters in this way confirmed the trends we found in our experiments with engineered EOD::PAR-3. First, as observed in EOD data, we observed a clear difference in diffusion behavior between smaller clusters (1mer, 2mer) and larger oligomers (3-4mer, >4mer) from the MSD analysis (Figure 6C-E). The anomalous parameters for 1mers and 2mers remain below 1 at all times, which indicate weakly confined diffusion, while the anomalous parameter for larger clusters increased from <1 on short time scales to >1 on longer time scales, which demonstrates that larger PAR-3 clusters display directed motions during the time scale of polarization. Second, as in EOD diffusion analysis, we calculated the diffusion coefficient, k_off_ and membrane binding half-time for each cluster size group. In agreement with the EOD data, we observed a decrease in the diffusion coefficient, a decrease in k_off_, and an increase in membrane half-time with increasing cluster size (Figure 6F-H).

Although we developed a systematic approach to estimate the cluster size, our analysis is inherently limited by incomplete labeling of the HaloTag with its ligand dyes. In previous work, we showed that in *C. elegans* zygotes, 70-80% of HaloTag molecules can be successfully labeled with ligand dyes (Dickinson et al., 2017; Sarıkaya and Dickinson, 2021). Since unlabeled molecules do not contribute to the fluorescence intensity of a cluster, incomplete labeling is expected to result in a systematic underestimation of cluster sizes in these experiments. With this in mind, two features of our dual labeling data merit further discussion.

First, we tracked many particles that had no detectable intensity in the abundant (far-red) channel and were therefore classified as PAR-3 monomers (Figure 6C). This is surprising because we did not observe any significant membrane stabilization of PAR-3 monomers in the PAR-3(RRKEEE) mutant strain (Figure 1; Dickinson et al., 2017). Some fraction of the PAR-3 population with estimated size of 1 are more likely to be 2mers in which the second molecule of PAR-3 failed to be detected. However, given the relatively large fraction of particles (672 out of 1,535 tracks) that appeared to be monomers, it is possible that these data may reveal a previously unappreciated ability of wild-type PAR-3 monomers to interact with the membrane (see Discussion).

Second, among PAR-3 particles estimated to be 2mers, we observed 2 populations that displayed different diffusive behavior. The majority of particles (284 out of 297 tracks classified as 2mers) underwent weakly confined diffusive motion, similar to the 1mers in this dual-labeling experiment and to engineered dimers. A small fraction (13 out of 297 tracks classified as 2mers) underwent directed motion on longer timescales, similar to larger oligomers. Considering the 70-80% labeling efficiency of the HaloTag in this system, this minority of ‘2mer’ tracks are very likely larger oligomers whose size was underestimated due to incomplete labeling.

Overall, these results are consistent with our analysis of engineered PAR-3 oligomers. We conclude that a size threshold of 3-4 monomers governs the diffusive behavior and advective transport of wild-type PAR-3 to establish the anterior-posterior axis of the *C. elegans* embryo.

## Discussion

Cortical flow is a general cell behavior that has been reported in a variety of cell types and serves several different functions (Ananthakrishnan and Ehrlicher, 2007). Here we have studied the biophysical basis for oligomerization-dependent protein transportation by cortical flow. We have used 2 novel methods – engineering PAR-3 size via EODs and dual-labeling of endogenous PAR-3 – to dissect the diffusive behavior of PAR-3 clusters of different sizes. Both approaches revealed a sharp decrease in diffusion coefficient and an increase in anomalous parameters as cluster size increases from two to four monomers, but showed only slight changes in diffusive behavior as clusters grow larger. These experiments indicate that the motion of smaller PAR-3 clusters is dominated by diffusion, while larger PAR-3 clusters are effectively transported by advective flow. These trends are consistent with our previous observations (Dickinson et al., 2017) but add an additional degree of quantitative precision and raise new questions about the physical and molecular basis for the sharp shift in diffusive parameters at the size threshold.

Our novel dual-labeling strategy can be widely applied for monitoring the dynamics of membrane proteins, as many as 35% of which may be oligomeric (Goodsell and Olson, 2000). Our approach was inspired by pioneering studies that achieved single-molecule labeling *in vivo* via low level transgenic expression of the target protein coupled with partial photo-bleaching to tune the levels of fluorescence (Robin et al., 2014), and by single-Molecule Speckle (SiMS) Microscopy via microinjection (Yamashiro and Watanabe, 2017). Dual HaloTag labeling is easier to carry out than these earlier approaches, and additionally allows tracking of sparse labels without sacrificing the ability to visualize the bulk cellular protein. This approach could be applied to other examples of protein oligomerization. For example, E-cadherin forms clusters on the cell membrane that have recently been shown to be crucially important for transmission of forces during tissue morphogenesis (Huebner et al., 2021). Although the dynamics of these clusters have been studied by bulk techniques such as FRAP, a dual-labeling technique could reveal turnover rate and diffusivity, potentially providing new insights into how E-cadherin clustering controls its behavior. As another example, receptor tyrosine kinases (RTKs) are known to be modulated by homo-oligomerization (Schlessinger, 2000), but how oligomerization controls RTK activity is still not understood in detail. A dual-labeling technique, coupled with activity reporters, could provide insight into this question. Finally, we anticipate applying dual-labeling to other oligomeric proteins within the PAR polarity system, including PAR-2 (Arata et al., 2016) and CHIN-1 (Sailer et al., 2015), to determine how clustering affects their dynamics on the plasma membrane.

PAR-3 is a peripheral membrane protein that associates with the lipid bilayer by binding to phosphoinositides (Horikoshi et al., 2011; Krahn et al., 2010; Wu et al., 2007). *In vitro,* the diffusion of phosphoinositide-binding proteins has been found to follow a remarkably simple relationship in which the diffusion coefficient decreases linearly with the reciprocal of the number of phospholipid molecules bound by each protein molecule (Knight et al., 2010; Ziemba and Falke, 2013). The differences in diffusivity that we measured for PAR-3 clusters of different sizes are on this same order of magnitude, consistent with the possibility that PAR-3 diffusivity is dominated by its lipid binding.

We and others have shown that monomeric mutants of PAR-3 do not bind stably to the plasma membrane (Figure 1) (Dickinson et al., 2017; Li et al., 2010a; Rodriguez et al., 2017), so it was surprising that we observed a large population of apparent monomers at the membrane in our dual-labeling experiment (Figure 6). Notably, in PAR-3 monomeric mutants, the oligomerization domain was disrupted by charge-reversal (RRKEEE) mutations (Dickinson et al. 2017) or by deleting the entire N-terminus (Li et al., 2010a; Rodriguez et al., 2017). One possible explanation for this discrepancy is that positively charged residues in the PAR-3 N-terminus, which are disrupted by the monomeric mutations, could directly interact with the negatively-charged surface of the lipid bilayer. Consistent with the presence of additional membrane binding interactions, wild-type PAR-3 particles had slower diffusivities and longer membrane lifetimes than engineered oligomers of the same sizes (compare Figure 5A-C to 6F-H). The isolated PAR-3 N-terminus does not bind the membrane on its own (Dickinson et al., 2017), but still could contribute to membrane targeting in the context of the full-length protein. Future experiments will explore the relationship between PAR-3 structure, oligomerization and membrane binding in greater detail.

In our previous study, the cell cycle kinase PLK-1 was shown to negatively regulate PAR-3 oligomerization by phosphorylating residues in the PAR-3 oligomerization domain (Dickinson et al., 2017). In that study, we attempted to isolate constitutive oligomerization mutants of PAR-3 by mutating the two identified PLK-1 target residues to alanine, but were unable to generate stable lines due to highly penetrant lethality and sterility. Consistent with this result, we were unable to isolate EOD::PAR-3 direct fusion lines through CRISPR-mediated insertion of EODs into the PAR-3 locus, which necessitated using the nAb-linking strategy instead. In light of these results, it was surprising that our EOD::nAb; YFP::PAR-3* strains not only are viable, but are healthier than the PAR-3* monomeric mutants. We do not have a clear explanation for this observation. One possibility is that PAR-3 monomers – which are expected to be absent from both the direct EOD::PAR-3* fusion strains and the PLK-1 phosphorylation site mutants, but present in the EOD::nAb; YFP::PAR-3* strains due to the difference in expression level between the EOD constructs and PAR-3 – might have a critical, unknown role in development. Alternatively or in addition, phosphorylation of PAR-3 by PLK-1 might regulate crucial PAR-3 interactions other than homoligomerization. These hypotheses represent interesting areas for further investigation.

Developmentally, the purpose of PAR-3 clustering is to transport the key polarity kinase, aPKC, to the anterior of the zygote, allowing asymmetric cell fate specification. PAR-3 oligomerization and its binding to PAR-6/aPKC are cooperative (Dickinson et al., 2017): PAR-3 clusters containing 3 or more subunits bound strongly to aPKC/PAR-6 in sc-SiMPull experiments, while dimers bound more weakly and monomers associated only at background levels (Dickinson et al., 2017). This observation was in agreement with earlier work showing that punctate PAR-3 and PAR-6/aPKC signals colocalize on the cortex (Beers and Kemphues, 2006; Dickinson et al., 2017; Hung and Kemphues, 1999; Lang and Munro, 2017; Li et al., 2010b). The threshold for PAR-3 binding to PAR-6/aPKC is strikingly consistent with our observed size threshold for PAR-3 clusters to be efficiently segregated by cortical flow, suggesting a possible connection between PAR-3 transportation and aPKC/PAR-6 binding. One hypothesis is that aPKC/PAR-6 binding could increase the volume of the aPAR complex, making it more susceptible to corralling effects and thus overcoming random Brownian diffusion. Alternatively, association with PAR-6/aPKC might facilitate membrane binding of PAR-3, either by inducing a PAR-3 conformational change (Chen et al., 2013) or through association with CDC-42 (Joberty et al., 2000) or other partners. Future work will attempt to test whether binding to PAR-6/aPKC is required for the shift in diffusive behavior that we report here.

## Supporting information

Key Resource Table

## Acknowledgments

We thank Luke Lavis for sharing JaneliaFluor dyes prior to publication, and Charles Lang, Edwin Munro, Jeanne Stachowiak, John Wallingford and members of the Dickinson lab for helpful discussions and comments on the manuscript. This work was supported by a Provost’s Graduate Education Fellowship from the University of Texas at Austin (YC), by a research grant from the Mallinckrodt foundation (DJD), and by NIH R00 GM115964 and R01 GM138443 (DJD). DJD is a CPRIT Scholar supported by the Cancer Prevention and Research Institute of Texas (RR170054). Some strains were provided by the Caenorhabditis Genetics Center, which is funded by the NIH Office of Research Infrastructure Programs [P40 OD010440].

## Author Contributions

YC and DJD designed the experiments. DJD designed and built the TIRF microscope used for dual-labeling experiments. YC performed the experiments. YC and DJD analyzed the data. YC carried out and analyzed the modeling studies. DJD supervised the project and secured funding. YC and DJD co-wrote the manuscript.

## Methods

### KEY RESOURCE TABLE

The Key Resource Table is provided as a supplemental file.

### CONTACT FOR REAGENT AND RESOURCE SHARING

Requests for resources and further information should be directed and will be fulfilled by the Lead Contact, Daniel J. Dickinson.

### EXPERIMENTAL MODEL AND SUBJECT DETAILS

All C. elegans strains were fed with OP50 and maintained on standard NGM growth medium. All strains were kept in a 20°C incubator unless noted otherwise. Embryos were examined before sex can be determined, however, most of embryos were likely to be hermaphrodites because the spontaneous occurance of male without mating is rare.

All genetic modifications to the C. elegans genome were made using protocols previously published by our laboratory (Dickinson et al., 2013, 2015). In brief, we used NEBuilder HiFi DNA Assembly to generate a plasmid construct that contains the homologous repair template with genome modifications and a selectable marker, flanked by 500-1500 bp of unmodified genomic homology arms. A repair template construct, a Cas9-sgRNA expressing vector, and a vector expressing extrachromosomal array fluorescent markers were co-injected into the syncytial gonads of young adult hermaphrodites. DNA repair by homologous recombination was triggered by Cas9 cleavage of the *C. elegans* genome, which allows the incorporation of modified repair template via homologous recombination. Knock-in animals were selected from the F2 progeny of injected animals using hygromycin selection and a phenotypic marker. After each knock-in strain was isolated, the selectable marker was excised by Cre-Lox recombination.

### METHOD DETAILS

#### TIRF Microscopy

Adult *C.elegans* carrying eggs were dissected in a drop of egg buffer on polylysine-coated coverslips. The embryos were gently flattened by mounting with 22.8 mm beads (Whitehouse scientific, Chester, UK) as spacers. Dual-labeling experiments (Figure 6) were carried out on a custom-built TIRF microscope (see below). All other TIRF images were acquired using a Nikon Eclipse Ti-2 microscope equipped with a 100X, 1.49 NA objective; a Photometrics Prime 95B camera; and an iLas2 circular TIRF illuminator (Roper scientific, É vry, France). TIRF images were magnified by a 1.5X tube lens before being collected by camera chip. The TIRF illuminator was operated in ellipse mode for acquiring images of whole embryos. mNG was excited using a 488 nm laser; YFP was excited using a 505 nm laser; mScarlet and JF585 were excited using a 561 nm laser; and JF646 was excited using a 638 nm laser.

#### Cortical Cluster Intensity Quantification

We used Utrack (Jaqaman et al., 2008) to detect the particle and to calculate the particle brightness in TIRF images. For particle detection, we used the point source detection algorithm with alpha = 0.01 and other parameters as default. In order to uncouple the effect on cluster size from cortical tension(Wang et al., 2017b), the movie was acquired using embryos dissected from MLC-4 RNAi treated worms (Figure 1E).

#### Cortical Flow Speed Quantification

Adult *C.elegans* carrying eggs were dissected in a drop of egg buffer on polylysine-coated coverslips. The embryos were gently flattened by mounting with 22.8 mm beads (Whitehouse scientific, Chester, UK) as spacers. DIC movies were acquired using a Nikon Eclipse Ti-2 microscope equipped with a 100X, 1.49 NA objective; a Photometrics Prime 95B camera, and Nomarski DIC optics. DIC images were magnified by a 1.5X tube lens before being collected by camera chip. One-cell stage embryos at establishment phase were imaged at 1 seconds per frame. Kymographs were generated by extracting a 20 pixel wide strip at the center of the imaged embryo image and stacked these strips on top of one another using the ‘‘montage’’ command in FIJI. The angle of a slope on the posterior was measured and the cortical flow speed in μm/min was calculated by:

Horizontal Displacement / min = tan(angle) X 60 frames X 20 pixels X 73.3 nm/pixel

#### Cell Polarization Quantification and Fluorescence Measurements

To quantify the polarization ability of our EOD strains, ovals matching the shape of the anterior half and posterior half of the embryo were drawn and the mean intensity per pixel was quantified using the Measure command in FIJI. For each embryo, the anterior intensity is normalized to 1 and the ratio of posterior intensity/anterior intensity was calculated.

#### Embryo Lethality and Sterility Quantification

A single young adult was picked to a new plate. After 12hrs, which allows the adult to lay eggs, the adult was removed from the plate. After another 24hrs during which the eggs were allowed to hatch, any unhatched embryos and hatched larvae were counted and embryonic lethality was calculated. The plate was maintained for additional 3 days for counting the number of fertile adults and sterile adults, which is used for calculating adult sterility.

#### Sc-SiMPull and Photobleaching Step Counting

We exactly followed the protocol in Dickinson et al., 2017. For reagents, experimental procedures, and data analysis software, please see Dickinson et al., 2017.

#### PAR-3 Particle Tracking and Motion Analysis

We used Utrack (Jaqaman et al., 2008) to track the motion of PAR-3 particles in TIRF images. For particle detection, we used the point source detection algorithm with alpha = 0.01 for EOD data and alpha = 0.03 for dual-labeling data, with other parameters as default. For the particle tracking step, we used Brownian + directed motion mode with minimum track length = 2 and maximum GAP to close = 3 for both EOD and dual-labeling experiments.

#### MSD analysis and Anomalous parameter visualization

Data from UTrack were converted to a suitable format using a custom MATLAB script, and then fed into the MSDanalyzer MATLAB function (Tarantino et al., 2014) to generate MSD plots. We eliminated tracks shorter than 10 frames (0.5 s of data), because we observed that UTrack sometimes erroneously identified short-lived particles in the noise of the images. To estimate slopes, we fit each averaged log-log MSD curve to a smoothing spline using the SLM toolbox for MATLAB. To avoid overfitting noisy data, the ends of each curve were trimmed prior to fitting.

Fits were constrained to be increasing functions and, where appropriate, concave-up. The slope of the curve is then estimated by taking the first derivative of the fitted spline.

#### Super-resolution iSIM Imaging

Adult *C.elegans* carrying eggs were dissected in a drop of egg buffer on polylysine-coated coverslips. The embryos were gently flattened by mounting with 22.8 mm beads (Whitehouse scientific, Chester, UK) as spacers. Images were acquired using a Nikon Eclipse Ti-2 microscope equipped with a 100X, 1.49 NA objective; a Photometrics Prime BSI camera; an OptoSpin filter wheel (CAIRN Research, Kent, England), and an vt-iSIM super-resolution confocal scan head (VisiTech international, Sunderland, UK). Confocal images were magnified by a 1.5X tube lens before being collected by camera chip. During image acquisition, the focal plane was centered at the cortex, and along with 2 slices above and below, 0.25 um per slice step, 5 total slices were collected. GFP was excited using a 488 nm laser and mScarlet was excited using a 561 nm laser.

#### Imaging Processing and Display

The high-resolution iSIM images (Figure 3C-D) were processed through FIJI to generate Z maximum projection images. The PAR-3 channel (Figure 3D) was processed using the RF denoise command in FIJI (theta = 10) after maximum projection. Following these operations, brightness and contrast were adjusted for visibility of the signals. No other image manipulations were performed.

To generate kymographs (Figures 3A), we extracted a 15 pixel wide strip at the center of the imaged embryo image and stacked these strips on top of one another using the ‘‘montage’’ command in FIJI.

#### Particle Image Velocimetry Quantification

To quantify the coupled motion of PAR-3 and actin filaments network (Figure S2B), we applied velocimetry (PIV) to the mSc::PAR-3 and GFP::Utrophin image channel using the PIVlab MATLAB plugin (Thielicke and Stamhuis, 2014). For mSc::PAR-3 channel: Images were pre-processed with the high-pass filter = 50, Wiener2 denoise filter = 5. The contrast was manually adjusted. Other preprocessing filters were disabled. Flow detection used the default FFT phase-space algorithm with 3 passes (window sizes 64, 32 and 16 pixels) and linear window deformation. Post-processing was done with velocity limits drawn manually to exclude outliers. Vectors that were rejected by these filters were replaced by interpolation. For Utrophin channel: Images were pre-processed with the high-pass filter = 400, Wiener2 denoise filter = 2. Other processing methods were the same as for mSc::PAR-3 channel.

#### Pearson’s Correlation

To quantify the correlation of velocity in PAR-3 and cortex (Figure S2C), the data was organized to a three dimensional matrix, with size of ( x-dimension, y-dimension, 2 (x-vector/y-vector)).

Pearson’s correlation was calculated by MATLAB Command: corrcoef (Utrophin velocity matrix, PAR-3 velocity matrix).

To quantify the colocalization of PAR-3 and actin, we imported the image sequence of mSc::PAR-3 and GFP::Utrophin into the matlab as a numerical matrix. Data was organized to a three dimensional matrix, with size of ( x-pixel position, y-pixel position, t-time resolution). Pearson’s correlation was calculated by MATLAB Command: corrcoef (Utrophin velocity matrix, PAR-3 velocity matrix).

#### RNA Interference

RNA interference targeting *mlc-4* to eliminate cortical flow was performed by injection.We amplified 1 kb of *mlc-4* from cDNA using primers that added T7 promoters to both ends of the amplified fragment. dsRNA was then synthesized using the T7 Ribomax kit (Promega) as instructed by the manufacturer. 1 mg/mL *mlc-4* dsRNA was injected into young adults in either GFP::Utrophin background or EOD;PAR-3 background, and the worms were dissected 24-28 hrs later for embryo imaging.

#### Calculating Diffusion Coefficients

The PAR-3 tracks information generated by Utrack was reformatted for the ‘msdanalyzer’ MATLAB function, and the results were processed by ‘msdanalyzer’ for generating MSD curves (Tarantino et al., 2014). The built-in ‘fitMeanMSD’ function of msdanalyzer was used to fit the first 6 time steps of each MSD curve to straight line, which has a slope of 4*D.

#### Membrane Half Life and k_off_ Quantification

To quantify the membrane half life of EOD::PAR-3 clusters (Figure 5E), we tracked YFP::PAR-3 clusters using Utrack. The Utrack output was translated into a more readable format using a custom-written MATLAB script. The membrane half was calculated from the dwell times of individual particles at the membrane, as follows (Kinz-Thompson et al., 2016):

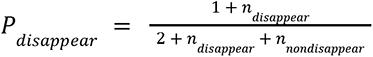

where the *n_disappear_* is the number of particle disappearance events, and *n_nondisappear_* is the sum of track length of all tracks;

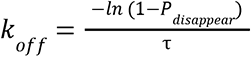

where τ is the measurement interval, which is 50ms for our experiments; and

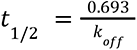

The 95% confidence intervals were calculated from the β distribution using the MATLAB function betaincinv (w, y, z, ‘lower/upper’). Where w = 0.025 for defining a 95% confidence interval, y = 1 + #of disappearing events(# of tracks), z = 1 + #of non-disappearing events.

#### Simulations of PAR-3 polarization

We developed a custom mathematical modeling script, written in MATLAB, for simulating PAR-3 cluster diffusion during polarization. The source code for our software is available at: https://github.com/IvyChang1994/Par3-Modeling.git

In brief, our simulation pipeline consists of the following steps: 1000 clusters with randomly distributed initial positions are generated. We simulated cluster positions for every time interval (0.05 second) during each 30-minute simulation.

1. Each cluster at each time point is moved by Brownian diffusion based on the displacement calculated from diffusion coefficient measured in the EOD live embryos (Figure 5A) and a direction generated randomly.
2. Each cluster at each time point is moved by cortical flow. According to previous studies, the cortical flow speed decreases almost linearly from posterior pole to the anterior pole at polarization stage, with the peak flow rate to be 7.7μm/min at posterior pole (Goehring et al., 2011a; Gross et al., 2019; Mayer et al., 2010).
3. Each cluster at each step has a probability of dropping off the membrane. We made the assumption that the number of clusters on the membrane remains constant because of the equilibrium of membrane binding and unbinding. So each cluster that drops off the membrane will re-bind at a new location determined by the current PAR-3 distribution. To generate the PAR-3 distribution at each time point, we counted the number of PAR-3 clusters in each 0.6μm bins. This distribution was smoothed using a moving average over 3 bins, and then new PAR-3 clusters were created at locations chosen at random from this smoothed distribution.

#### Polarity state quantification for simulated embryos

To characterize the polarization state of a simulated embryo (Figure 5D), we calculated the percentage of clusters present in the anterior domain as a function of time. The anterior domain is defined as the left side of the cleavage furrow, which is 57% percent of the embryo length (Dickinson et al., 2017). There are 36,000 time intervals in each simulated segregation, which causes sharp fluctuations. We therefore smooth the data before plotting using MATLAB function ‘smooth’, with smooth range set to 3600. The average at each time point across 10 simulations were calculated for generating the black average line.

As a second approach (Figure 5E), we extracted the x coordinates from final cluster positions. The coordinates were binned into 6μm x-axis ranges and the number of clusters in each bin were counted.

#### HaloTag Ligand Dye Feeding and Dual-labeling Experiments

JaneliaFluor Dyes (gifts from Luke Lavis) were dissolved in acetonitrile to 1 mM, dispensed into 2 μL aliquots in PCR tubes, dried under vacuum and stored at -20C in a desiccator upon receipt. Before use, 2 μL of DMSO was used to dissolve each single-use aliquot. 1 mL of OP50 liquid culture was spun down and resuspended in 100 μL S medium (150 mM NaCl, 1 g/L K2HPO4, 6 g/L KH2PO4, 5 μg/L cholesterol, 10 mM potassium citrate pH 6.0, 3 mM CaCl2, 3 mM MgCl2, 65 μM EDTA, 25 μM FeSO4, 10 μM MnCl2, 10 μM ZnSO4, 1 μM CuSO4) for each dye. Dissolved dye was added to the S medium-bacteria mixture. For the dual-labeling experiment, 2 μL of JF585-medium-bateria mix was added to 100 μL JF646-medium-bateria mix. For the double dilution experiment, a separate 100 μL medium-bacteria mix was made, and 2 μL of JF585-medium-bateria mix and 1 μL of JF646-medium-bateria mix was added to the 100 μL medium-bacteria mix. 30 µL of dye(s)-medium-bateria mix was dispensed into wells of a 96-well plate with round-bottom wells, and 20-30 L4-stage worms were picked into each well. Worms were grown at 20°C, with shaking at 230 rpm, for 16-24hrs before imaging. The final JaneliaFluor dye concentration used for feeding was 15uM.

#### Micromirror TIRF Imaging

Adult C.elegans carrying eggs were dissected in a drop of egg buffer on polylysine-coated coverslips. The embryos were gently flattened by mounting with 22.8 mm beads (Whitehouse scientific, Chester, UK) as spacers. TIRF images were acquired using a custom-built TIRF microscope equipped with an Olympus APON 60X, 1.49 NA objective; a Photometrics Prime 95B camera; and a dual-view emission path design. This microscope uses a micromirror illumination path design that is optimal for imaging multiple wavelengths simultaneously (Friedman and Gelles, 2015; Friedman et al., 2006). After collection of fluorescence emission by the objective lens, an image was formed using a tube lens (Edmund Optics) with a 400 mm focal length, resulting in 133.33X magnification. The red and far-red channels were split using a T635lpxr dichroic mirror (Chroma), passed through a set of relay lenses, and collected side-by-side on the camera chip. The relay lenses introduced an additional 1.2X magnification, so that the total system magnification was 160X. JF585 and JF646 were simultaneously excited using 50mW 561 nm laser and 50mW 637 nm laser.

#### Dual-labeling Experiments Data Analysis

We calibrated JF646 intensity using control experiments in which both JF585 and JF646 were diluted to single-molecule levels. As expected, the measured fluorescence intensities of single dye molecules followed a log-normal distribution (Figure S4A-B) (Mutch et al., 2007). The measured single-fluorophore intensities varied from embryo to embryo, which is not surprising and likely reflects variations in mounting and eggshell thickness; importantly, however, the intensities of the red and far-red channels varied in tandem (Figure S4A). We therefore adopted the following calibration procedure, in which we used the red dye intensity as an internal standard, since it is present at single-molecule levels in both sets of experiments. First, we normalized our double-dilution datasets to the red dye intensity. Normalization was done in log space, to account for the log-normal shape of the intensity distribution, and was effective at correcting for embryo-to-embryo variations in fluorescence intensity (Figure S4A-C). Second, we calculated the ratio of the average far-red dye intensity to the average red dye intensity (Figure S4D). Third, the intensity of red dyes in each sparse/abundant dual-labeling experiment was measured, and the intensities were normalized to the mean red dye intensity to allow comparison between embryos (Figure S4E-H). The cluster size information of PAR-3 labeled by red (JF585) fluorophore was acquired by a custom-written MATLAB script. This script draws a box in the far-red channel at the location where the red signal is located, and calculates the fluorescence intensity in the far-red channel by subtracting local background from a larger box. Fourth, from the calibrated red/far-red intensity ratio, an expected mean far-red intensity per molecule was calculated. The source code for our software is available at https://github.com/IvyChang1994/getClusterSize.git.Finally, this calculated far-red intensity per molecule was used to estimate the number of far-red labeled Halo::PAR-3 molecules in each tracked cluster. This calibration procedure accounts for the differences in global fluorescence intensity between embryos, presumably caused by the differences in laser penetration depth and eggshell thickness, and allows us to estimate the number of PAR-3 molecules per cluster.

---

**Figure S1.**
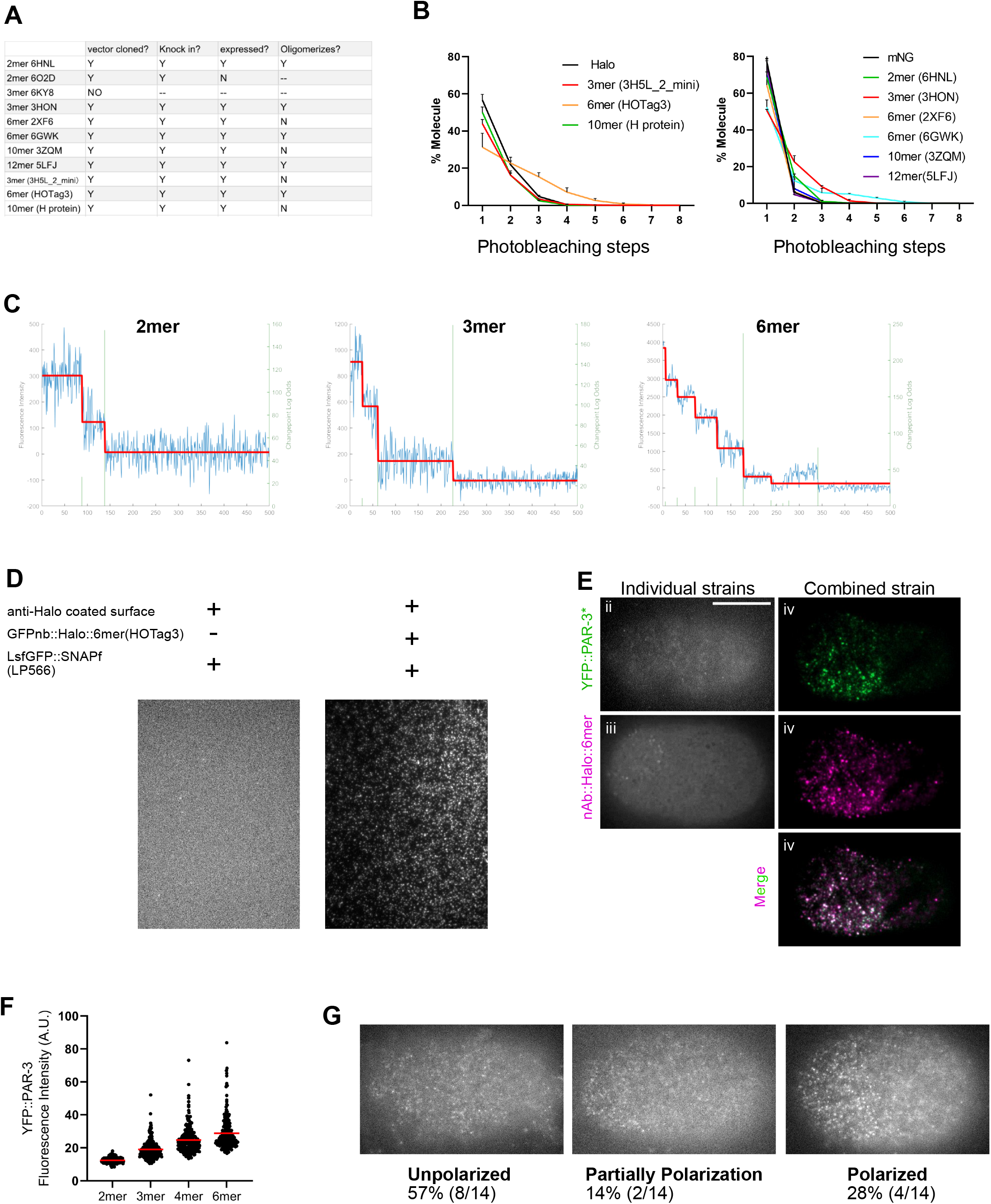
(A) Summary of the potential EOD domains tested. (B) The validation of EOD oligomer sizes using sc-SiMpull followed by photobleaching step counting. (3 replicates for each strain). EODs attached to Halo and tested by photobleaching Halo (left panel) and EODs attached to mNG and tested by photobleaching mNG (right panel). Please note that under this methodology, the photobleaching steps are predicted to be smaller than the real number of molecules in an oligomer due to the non-perfect maturation rate (50-80%) of fluorescent proteins and simultaneous bleaching of multiple fluorophores. (C) Examples of photobleaching traces of 2mer, 3mer and 6mer bleaching in 2 steps, 3 steps and 6 steps respectively. (D) sc-SiMpull assay testing the binding between GFP nanobody and GFP. The device surface was functionalized with anti-Halo antibodies. First, a GFPnb::Halo::6mer embryo was lysed to capture the EOD construct by means of the Halo – anti-Halo interaction. Then, a second embryo expressing sfGFP::SNAPf embryo was lysed to test whether the GFP nanobody, present in the EOD construct, could capture sfGFP. TIRF imaging for GFP signal. (E) Detailed illustration of the rescue experiment result, hexamer(HOTag3) was shown as an example. Left panels: (Top) TIRF imaging of embryo from YFP::PAR-3* monomeric mutant strain after polarization stage, corresponding to ii in panel figure 1A. (Bottom) TIRF imaging of embryos from GFPnanobody::HaloTag::6mer(HOTag3) construct strain after polarization stage, corresponding to iii in panel figure 1A. Right panels: TIFR imaging of polarized zygote from YFP::PAR-3;nAb::HaloTag::6mer(HOTag3) strain, corresponding to iv in panel figure 1A (Top) YFP::PAR-3 channel. (Middle) nb::Halo::6mer(HOTag3) channel. (Bottom) composite image with both YFP and Halo signal. Scale bar: 10 μm. (F) YFP fluorescence intensity of the foci on the cortex of YFP::PAR-3*;nAB::BFP::EOD embryos dissected from *mlc-4* RNAi treated worms. (G) The degree of polarity defect in the 2mer strain and the fraction of each category.

**Figure S2.**
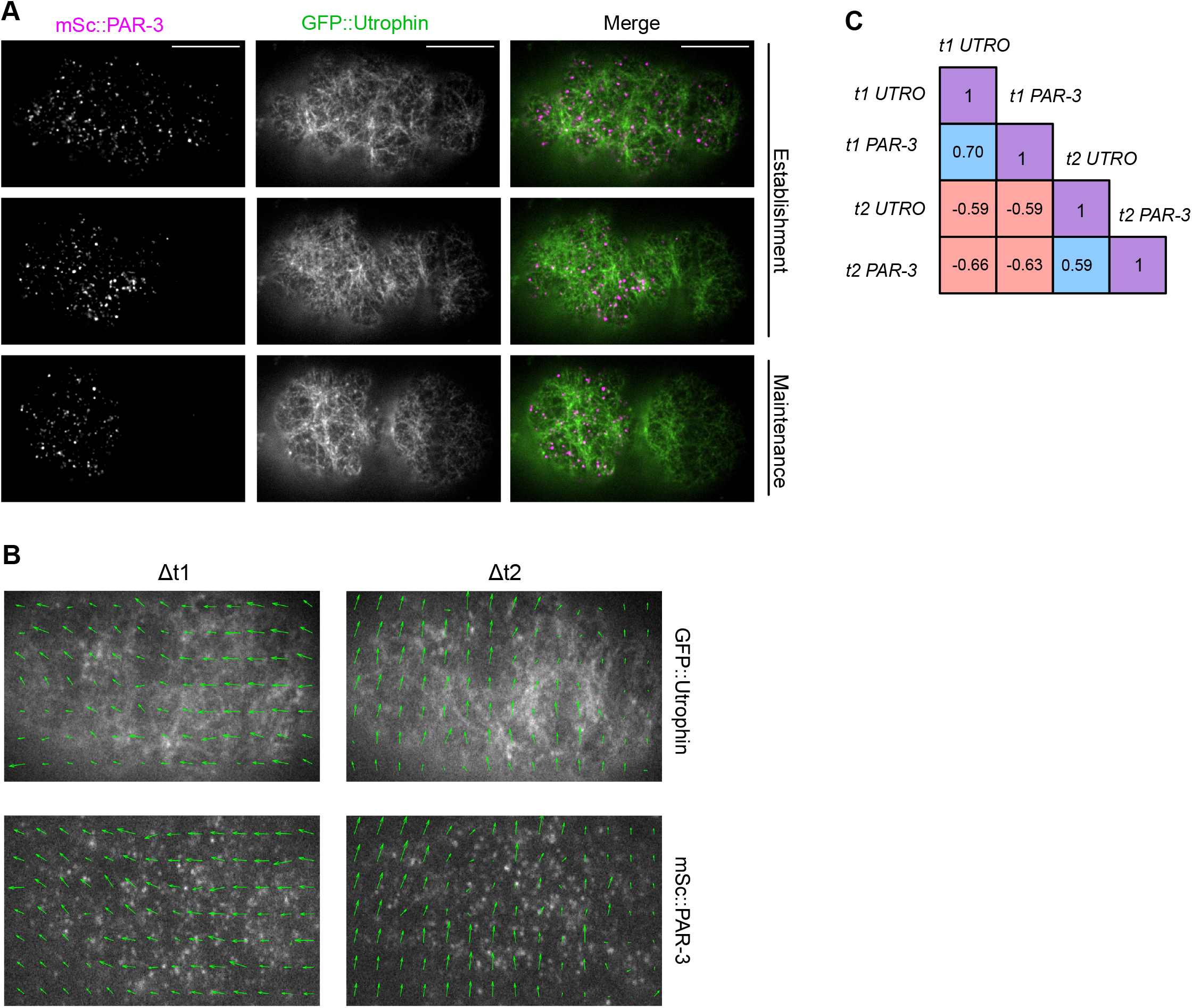
(A) TIRF imaging of mSc::PAR-3;GFP::Utrophin zygote. Anterior to the left. Scale bar = 10µm. (B) The vector field generated by PIVlab at the polarization stage. (C) Pearson’s correlation of the vector fields in (B). Blue boxes represent experiments where the vector field of actin and PAR-3 from the same time point is compared, red boxes represent negative controls where unrelated images are compared, and purple boxes serve as positive controls where the same image is compared to itself.

**Figure S3.**
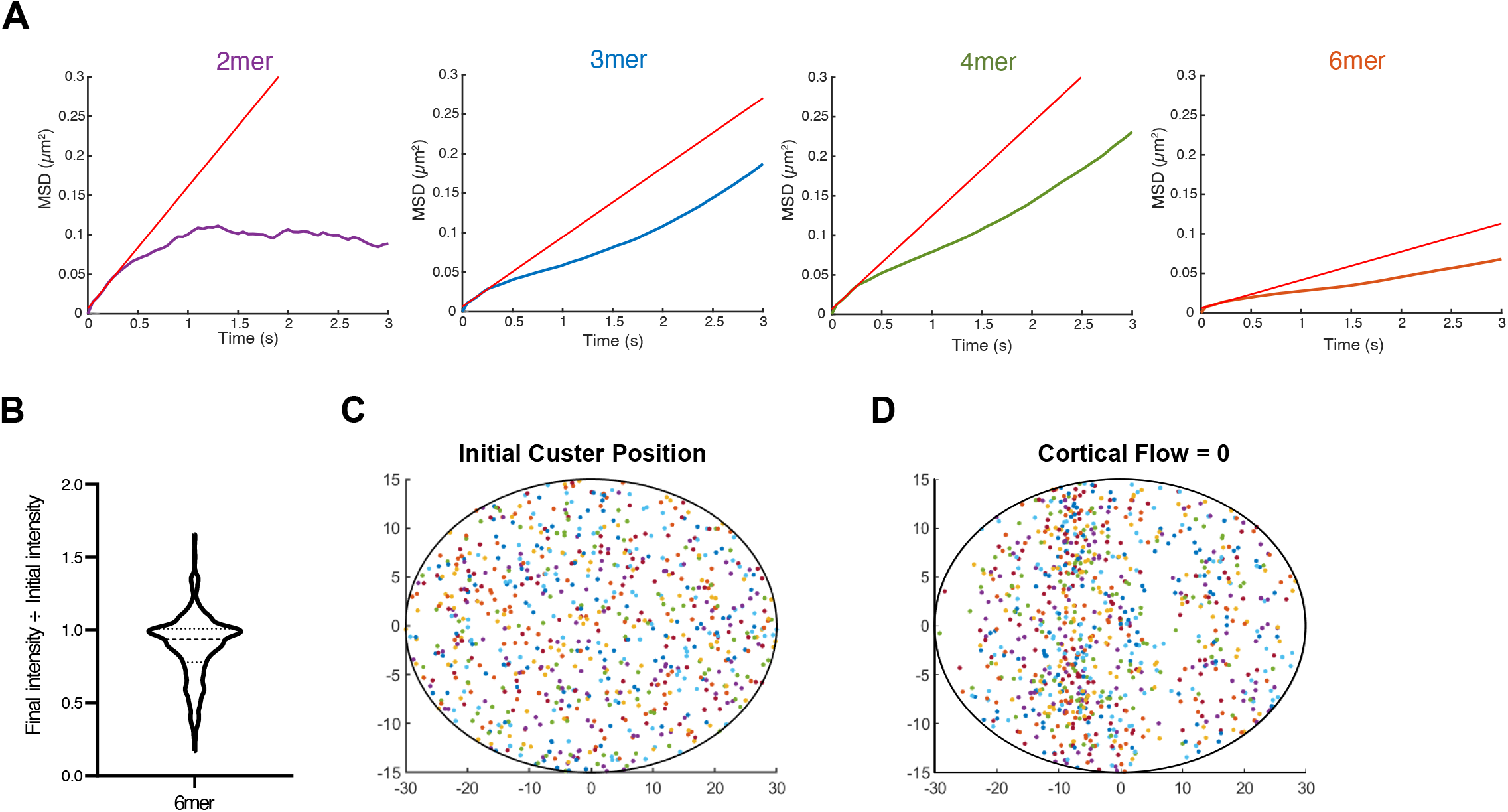
(A) Curve fits used to determine the diffusion coefficient for engineered PAR-3 oligomers. (B) Ratio of final:initial cluster intensity for 6mers under our imaging conditions. n=4,309 particles. (C) An example of the random initial cluster positions used for simulations. (D) PAR-3 cluster segregation simulated with cortical flow = 0 globally. Positive feedback leads to transient enrichment of particles in random locations but does not result in polarity.

**Figure S4.**
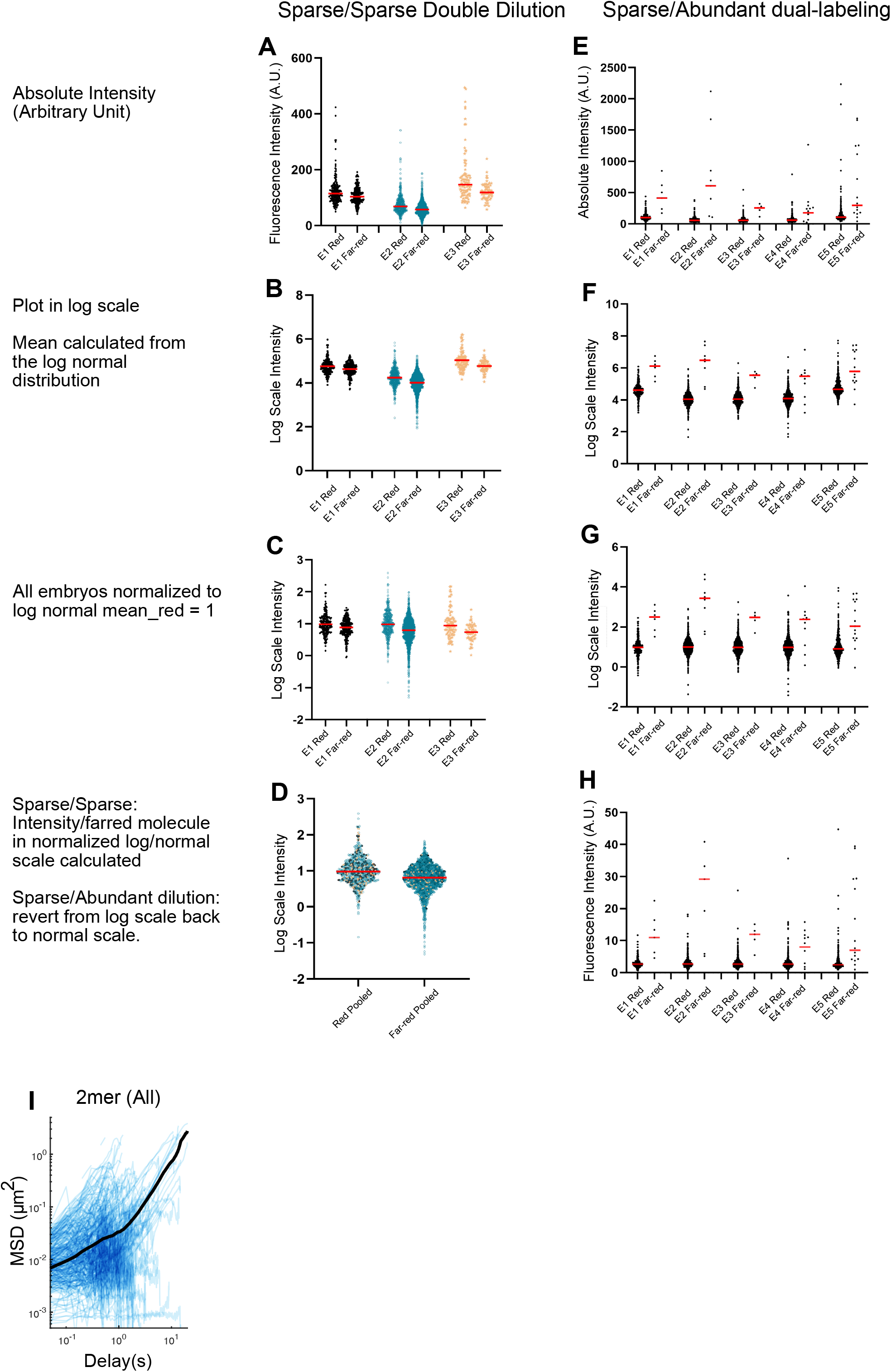
(A) The global intensity of red and farred signals in double-dilution experiments. Each dot represents a single foci detected. Red lines indicate the means. (B) The log scale intensity of (A). (C) The datasets in (B) are normalized to 1. (D) All 3 datasets from the same color channel are pooled together for calculating the single red/single far-red ratio. (E) The global intensity of red and the farred channel intensity of particles of interest in sparse/abundant dual-labeling experiments. Each dot represents a single foci detected. Red lines indicate the means. (F) The log scale intensity of (E). (G) The datasets in (F) are normalized to 1. (H) The normalized log scale intensity is converted back to normal scale, ready for converting into cluster size. (I) log/log scale MSD curves for all 2mers, each curve describing the motion of a single PAR-3 cluster. Data was acquired and pooled from 3 embryos.

## Movie Legends

Movie S1. TIRF live imaging of polarizing zygotes from strains with genetic modifications that result in different PAR-3 cluster size. Imaged at 3 seconds/frame. Anterior to the left.

Movie S2. iSIM live imaging of polarizing mNG::PAR-3; NMY-2::mKate2 zygote. Imaged at 3 seconds/frame. Anterior to the left.

Movie S3. TIRF live imaging of polarizing mSc::PAR-3; GFP::utrophin zygote. Imaged at 3 seconds/frame. Anterior to the left.

